# Does the Inertia of a Cell Depend Upon its Information-Content?

**DOI:** 10.1101/2020.09.09.285106

**Authors:** Ahmed Elewa

**Author notes:** e t: @egypsci.

## Abstract

A time dilation factor is derived from information entropy via the holographic principle. By imagining the embryonic cell as a hologram, time dilation during embryogenesis is predicted from the entropy of active genetic information and shown to be consistent with observed lengths of cell duration. The connection between entropy and cell duration as well as the corollary of increase in cellular mass with the reduction in information entropy are discussed. This preprint is the second in a series exploring the value of *Developmental Spacetime* as a framework for falsifiable predictions about the universe.

## INFORMATION ENTROPY AND TIME DILATION

The holographic principle states that the maximum entropy *N* of a closed volume is equal to the surface area divided by the smallest unit of area (Planck’s area *l*_P_ ^2^)(1).

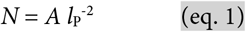

Since Planck’s length *l*_P_ = (*ħ G c*^−3^)^1/2^, time *t*_P_ = (*ħ G c*^−5^)^1/2^ and mass *m*_P_ = (*ħ c G*^−1^)^1/2^ are all expressed using the same constants *ħ*, *G* and *c*, it is possible to express Planck’s time as *t*_P_ = *l*_P_ *c*^−1^. From (eq. 1), we can also express Planck’s time as *t*_P_ = (*A N*_P_^−1^)^1/2^ *c*^−1^. *N*_P_ refers to the number of bits needed to represent an equiprobable Planck-scale system (i.e., maximum entropy). However, if the entropy of a system is no longer maximal and some components are represented more than others, a value lesser than *N*_P_ in quantity represented by Shannon’s entropy *H*_P_ would represent the information within the system (2).

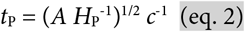

From (eq. 2), one can calculate Planck’s time for a system with entropy *H*_P_. However, since it is untenable to calculate the entropy of an ideal Planck-scale system with an enormous number of units, one can instead calculate the entropy of a representative subset of the system as follows. From (eq. 1) and the definition of *l*_P_, we can express *A* as *N*_P_ *ħ G c*^−3^ and write (eq. 2) as *t*_P_ = (*N*_P_ *ħ G c*^−3^ *H*_P_^−1^)^1/2^ *c*^−1^. This step relieves the equation from the explicit declaration of area since it is still conveyed in the number of bits *N*_P_. Next, to generalize the equation so that it is applicable to non-Planck scale holograms, we can relate the calculated entropy of that system to *N*_P_ by normalizing *H* to the maximum entropy of the system in question and multiplying that ratio by *N*_P_ (i.e. *H N*^−1^ *N*_P_), leading to *t*_P_ = (*N*_P_ *ħ G c*^−3^ *H*^−1^ *N N*_P_^−1^)^1/2^ *c*^−1^.

Simplification cancels out *N*_P_ altogether and recovers Planck’s time from (*ħ G c*^−5^)^1/2^ resulting in *t*_P_ = *t*_P_ (*N H*^−1^)^1/2^, which describes the dilation that occurs to Planck’s time due to the entropy of a hologram and can be generalized to time in general as

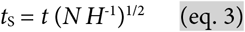

where *t*_S_ is Shannon’s time, or time with the effect of Shannon’s entropy taken into consideration. Deriving *t*_S_ predicts a dilation in time coinciding with a reduction in Shannon’s entropy. The following section evaluates this prediction using the cell as a hologram and gene expression as a proxy for information entropy.

## CEULLULAR ENTROPIC TIME DILATION

Nematodes, in particular *C. elegans*, have served as important models for developmental biology (3). A combination of small size (~1 mm long), transparency, and a limited number of cells (~ 1000) has enabled researchers to the definitively map the origin and fate of each cell in *C. elegans*, from a single cell to a mature adult (4) (see complete lineage maps at wormatlas.org/celllineages.html). To an external observer, the time an embryonic cell spends before dividing increases with each division (**Figures 1A & 1B**). The increase in cell duration can be expressed as 1/(1−*k*), where *k* the slope of cell duration increase with developmental time (5) (**Figure 1C**). For example, in the C cell lineage that eventually gives rise to the pharyngeal muscle cells, a slope of *k* = 0.23 captures the rate of increase in cell duration whereby the duration of a cell is approximately equal to 1/(1−*k*) multiplied by the duration of its mother (**Figure 1D**).

**Figure 1.**
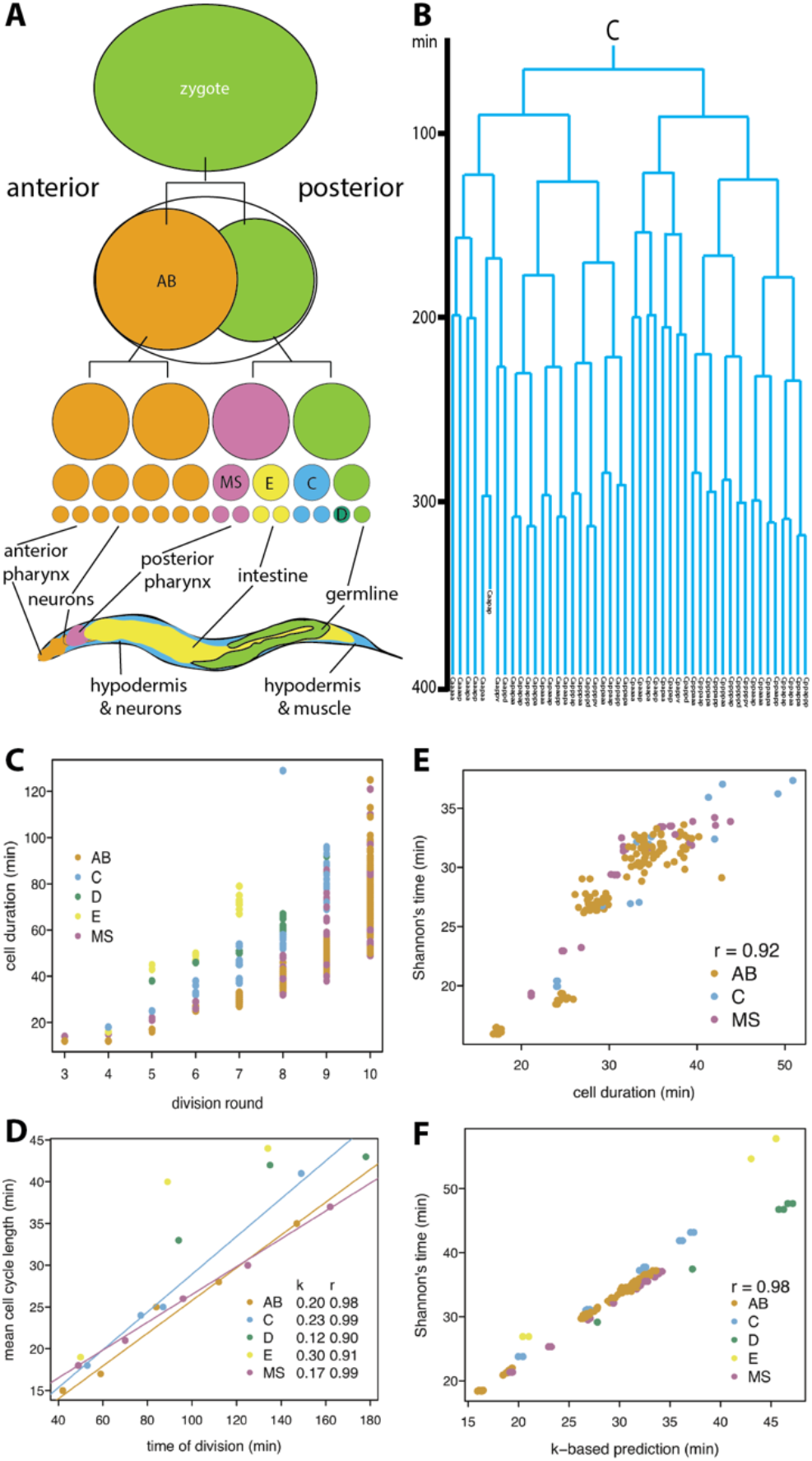
Predicting cell duration in the *C. elegans* embryo. **A)** A brief outline of the first four rounds of embryonic division and the body parts built by the AB, MS, E and C lineages. **B)** The lineage of the C blastomere as an example of increase in cell duration with each round of division. Each horizontal branch depicts a cell division and length of the vertical lines convey time. **C)** The gradual increase in cell duration during the first ten rounds of division based on (8). The five lineages are depicted in different colors per legend. **D)** Cell duration follows a geometric progression that differs per lineage. The inset lists the linear fitting for all lineages, *k* and their correlation coefficient *r*. **E)** Shannon’s time prediction correlates with observed cell duration (*p*-value <10^−15^). Minor lineages E and D are excluded from plot. **F)** Shannon’s time prediction correlates with *k*-based predictions (*p*-value <10^−15^).

Within each embryonic cell is a dynamic transcriptome, a set of RNA transcripts reflecting gene activity. The Shannon’s entropy of a cell’s transcriptome can be calculated from single-cell RNA sequencing data. Within the limits of present technology, the single-cell transcriptome is a representative sample of cellular information entropy. Therefore, from (eq. 3), the duration of a cell is equal to that of its mother after incorporating the effect of entropy

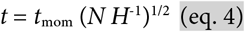

where *t* and *t*_mom_ are the durations of a cell and its mother, respectively, according to an external observer, *N* is the logarithm of the *number of expressed genes* (i.e. maximum entropy) and *H* is Shannon’s entropy calculated from the single-cell transcriptome (6, 7). Applying (eq. 4) to the first 8 out of 10 rounds of embryonic cell division in *C. elegans* (5) demonstrates that the dilation factor (*N H*^−1^)^1/2^ can predict cell duration (**Figure 1E**, Pearson’s correlation 0.92 and 0.85 when including D and E lineage). Extending the analysis to an independent set of cell durations observed at 20°C and 22°C degrees and including the ninth and tenth rounds of division (8) further demonstrates the utility of the proposed dilation factor (Pearson’s correlation 0.88 at 20°C and 0.87 at 22°C **Supplementary Table and Methods**). On average, predictions made using (eq. 4) correlated with those made with *k* (**Figure 1F**, Pearson’s correlation 0.98 **Supplementary Table and Methods**) but were ~10% shorter than observed cell durations (**see Discussion**). In conclusion, by considering the *C. elegans* embryonic cell as a hologram and its transcriptome a proxy for information entropy, *t*_S_ = *t* (*N H*^−1^)^1/2^ predicts time dilation after considering the effect of information entropy.

## DISCUSSION

In addition to *C. elegans*, the fruit fly (*Drosophila melanogaster*) and the African clawed frog (*Xenopus sp.*) have informed much of our understanding of embryonic development. *Drosophila* embryos proceed through thirteen rounds of synchronous cell divisions each lasting less than 10 minutes. Similarly, *Xenopus* embryos undergo twelve rounds of cell cleavage that are more or less similar in duration (9). Thus, early *Drosophila* and *Xenopus* embryogenesis do not display the same geometric progression seen in *C. elegans*. While this may seem to undermine the proposed connection between information entropy and cell duration, a key difference between *Drosophila* and *Xenopus* on one hand and *C. elegans* on the other must be considered. Fly and frog embryos depend on a deposit of maternal transcriptome during the initial synchronous cell divisions and shift to a reliance on their own genetic material afterwards. While these early embryonic cells differentially segregate their maternal information (and may therefore differ in transcriptome profiles), such reshuffling is unlikely to have a significant impact on overall information entropy. *C. elegans* embryos, however, shift to a dependence on their own genetic material after the second round of divisions and with the influx of newly synthesized transcripts, information entropy decreases as cells differentiate. Indeed, once *Xenopus* embryos transition to a dependence on their own gene expression a geometric progression in cell duration is observed (10). Another key difference between *Drosophila* and *C. elegans* embryos, is that the former is not composed of independent cells but instead is a syncytium. In other words, the cell in an early *Drosophila* embryo is not the closed volume envisaged in the holographic principle (eq. 1) and may therefore not be a suitable model to test the derived time dilation factor.

A reduction in transcriptome entropy corresponds with cellular differentiation, the process whereby a cell refines its identity and function and consequently the set of expressed genes and the magnitude of their expression. A differentiated cell expresses fewer genes and expresses them at higher [normalized] rates compared to a more pluripotent cell, which expresses more genes and expresses them at lower [normalized] rates (11, 12). Therefore, to a biologist the results described here may be trivial, for the rate of cell proliferation tends to decrease as the embryo differentiates and cells take on their terminal functions. The novelty here, however, is linking this cellular behavior to information entropy in a general manner consistent with non-biological systems including black holes; a predecessor to the holographic principle. A fact that is trivial to a biologist is the reduction in developmental rate with the reduction in temperature (within the limits of ectothermal physiology). For example, *C. elegans* develops at a slower rate when reared at 18°C compared to 23°C. The results described here propose a parallelism between thermodynamic and information entropy and their relation to physiological time.

Predicted cell durations based on (eq. 4) were generally ~10% shorter than observed durations. This deficiency could be a result of limiting sequencing and gene expression analysis to protein-coding genes and not accounting for non-coding genes and different alternative splice isoforms. Such limitations would necessarily affect the calculated information entropy.

Finally, a similar process leading to (eq. 3) can be carried out using Planck’s mass instead of time to yield Shannon’s mass, *m*_S_ = *m* (*N H*^−1^)^1/2^, or mass with the effect of Shannon’s entropy taken into consideration. This predicts an increase in the mass of a cell in the *C. elegans* embryo proportional to its increase in duration. More generally, this corollary predicts that a geometric progression in cell duration coincides with a similar increase in cell mass. Biologically speaking, this increase in mass may correspond to a significant increase in transcription and other forms of biosynthesis to fulfill cellular roles determined by differentiation. Two immediate considerations are the source of the increasing mass and the mass of terminally differentiated cells (i.e. with maximum cell duration). The embryo consists of extracellular spaces that can be endowed with building blocks for intracellular synthesis. For example, the fluid-filled cavity of the early *Xenopus* embryo can provide the matter required for increasing cellular mass as information entropy decreases so that while the overall mass of the embryo may remain constant during differentiation, the inertia of individual cells may increase. A similar extracellular source can be invoked in the case of *C. elegans*, for while the embryo is far more compact, extracellular spaces between the cells and eggshell do exist. With regards to terminally differentiated cells, consider a mother cell that lasts for two hours before dividing into daughter cells that last for two weeks. Will the post-embryonic daughters be 168 times more massive than their mother? A total accounting of biosynthesis may indeed confirm this hypothesis. The fact that final mass of the daughter cells may not differ much from their mother can be accounted for by energy-dependent degradation of bioproducts.

The gradual increase in Shannon’s mass with differentiation allows this quantity to replace epigenetic tension as a basis for the curvature of a modeled Waddington Epigenetic Landscape (13). Generating a Waddington Epigenetic Landscape using Shannon’s mass is a step towards articulating *Developmental Spacetime*; a conceptual and physical connection between expressed genetic information, cell time and mass and organismal geometry.

## SUPPLEMENTARY TABLE

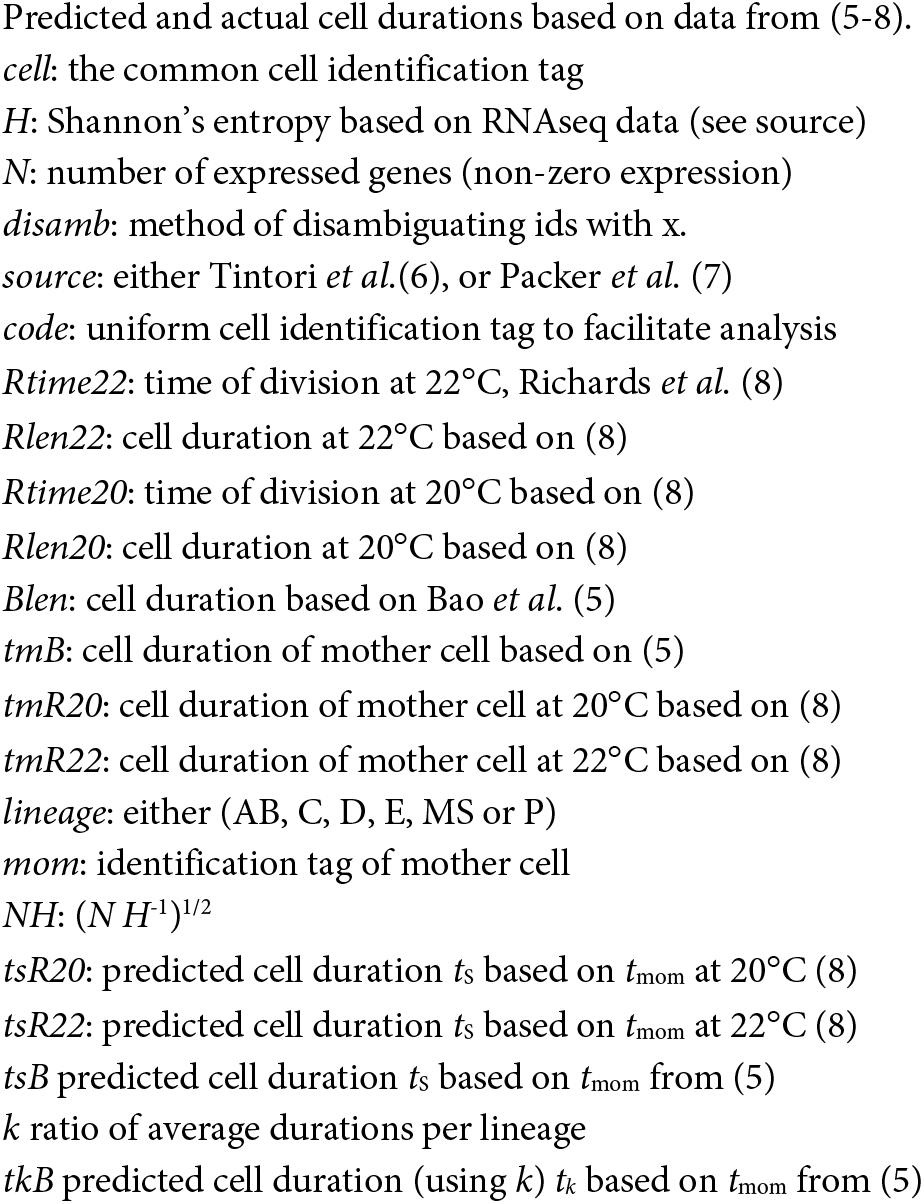

## METHODS

<u>Calculating information entropy</u> from single-cell RNAseq data (6, 7) was done as follows. For each cell, Shannon’s entropy *H* was calculated in R as-sum(1*p*\*log(*p*,2)) where *p* is the bootstrap median transcripts per million (normalized to 1) in case of (7) or the mean reads per kilobase of transcript (rpkm) of replicates in case of (6). *N* in eq. 4 was defined as the number of genes with non-zero gene expression. For (6), four ambiguous cells (MSx1, MSx2, Cx1, Cx2) were converted to MSa, MSp, Ca, and Cp, respectively. The values for ambiguous cells ABalx, ABarx, ABplx and ABprx were each duplicated and assigned to separate cells where x was either anterior (a) or posterior (p). The dataset in (7) consisted of over 500 cells many of which included of at least one ambiguous cell (an x instead of a definite division axis a/p, l/r or d/v). For such ambiguous cells, entropy was calculated and then assigned to two cells covering the possible assignees (e.g. Cxap to Caap and Cpap). In the end the final dataset (**Supplementary Table**) consisted of 458, 87, 49, 18, and 15 cells for AB, MS, C, E and D lineages, respectively. <u>Cell duration</u> was based on (5, 8). In the case of (8), cell duration at 22°C was obtained by subtracting the time of division from that of its mother. <u>Calculating *k*</u> was done in accordance with (5). For each lineage the average cell duration (cell cycle length) for each division round was compared to the average time of division of that round. Line fitting was performed in R using line() and the resulting coefficients were used to define intercept and slope (*k*). <u>Statistical significance</u> (*p*-value) of all reported Pearson’s correlations was below 10^−15^ per cor.test() in R. Moreover, significance was visualized by randomizing the predicted values 1000 times and plotting histograms of correlations. Histograms were either based on all cells reported in **Supplementary Table**(below) or after removing duplicated values due to cell ambiguity (not shown).

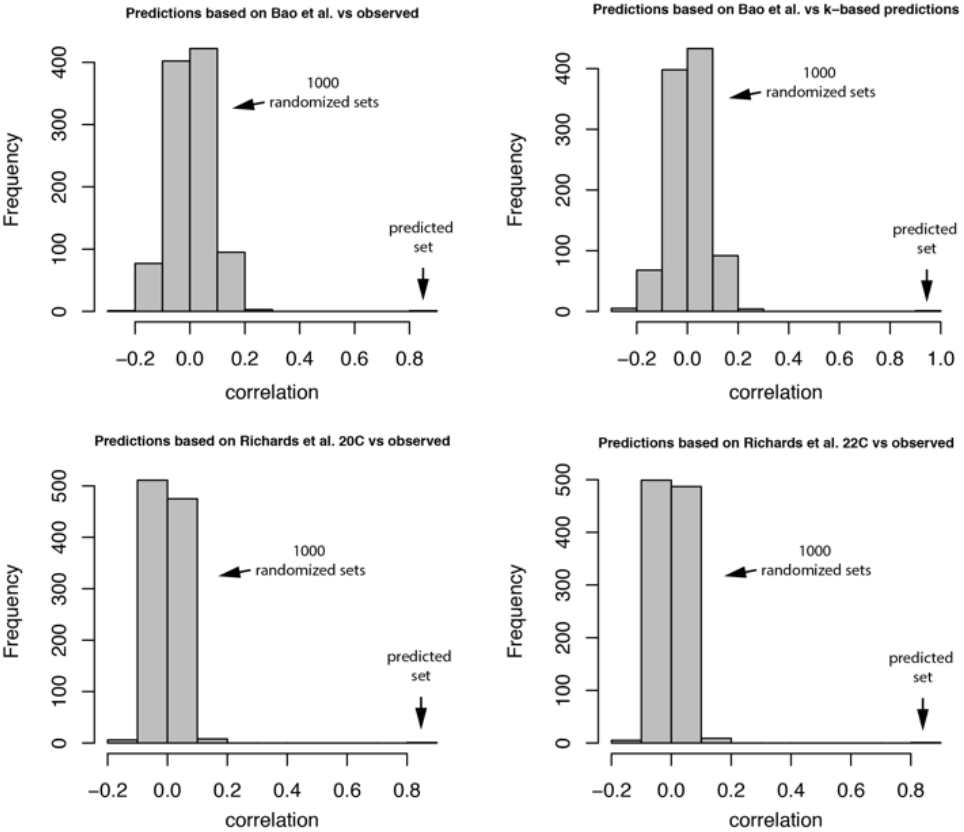

## ACKNOWLEDGEMENT

This work was made possible by the patronage of Mostafa Nashaat.

## REFERENCES

1. G. ‘t Hooft, The Holographic Principle. arXiv:hep-th/0003004 (2000). DOI: 10.1142/9789812811585_0005

2. C. Shannon, A mathematical theory of communication. Bell System Technical Journal 27, 379–423, 623–666 (1948).

3. S. Brenner, The genetics of *Caenorhabditis elegans*. Genetics 77, 71–94 (1974).

4. J. E. Sulston, et al., The embryonic cell lineage of the nematode *Caenorhabditis elegans*. Dev Biol 100, 64–119 (1983).

5. Z. Bao, et al., Control of cell cycle timing during *C. elegans* embryogenesis. Dev Biol 318, 65–72 (2008).

6. S. C. Tintori, et al., A Transcriptional Lineage of the Early *C. elegans* Embryo. Dev Cell 38, 430–444 (2016).

7. J. S. Packer et al., A lineage-resolved molecular atlas of *C. elegans* embryogenesis at single cell resolution. Science 365(6459) (2019).

8. J. L. Richards, et al., A quantitative model of normal *Caenorhabditis elegans* embryogenesis and its disruption after stress. Dev Biol 374, 12–23 (2013).

9. J. Newport, M. Kirschner, A major developmental transition in early *Xenopus* embryos: I. Cell 30, 675–686 (1982).

10. J. Newport, M. Kirschner, A major developmental transition in early *Xenopus* embryos: II. Cell 30, 687–696 (1982).

11. M. Percharde et al., Hypertranscription in Development, Stem Cells, and Regeneration. Dev Cell 40, 9–21 (2017).

12. S. Efroni et al., Global transcription in pluripotent embryonic stem cells. Cell Stem Cell 2, 437–447 (2008).

13. A. Elewa, A Waddington Epigenetic Landscape for the *C. elegans* embryo. bioRxiv, (2020). DOI: 10.1101/2020.01.01.892414

