## Supplementary Table for "Does the Inertia of a Cell Depend Upon its Information-Content?"

| cell | H | N | disamb | source | code | Rtime22 | Rlen22 | Rtime20 | Rlen20 | Blen | tmB | tmR20 | tmR22 | lineage | mom | NH | tsR20 | tsR22 | tsB | k | tkB |
|---|---|---|---|---|---|---|---|---|---|---|---|---|---|---|---|---|---|---|---|---|---|
| ABa | 11.8785 | 8332 | manual | Tintori | Xaa | 42 | NA | 42 | 12 | NA | NA | NA | NA | AB | Xa | 1.0471 | NA | NA | NA | 0.1962 | NA |
| ABp | 11.8436 | 8093 | manual | Tintori | Xap | 42 | NA | 42 | 12 | NA | NA | NA | NA | AB | Xa | 1.0470 | NA | NA | NA | 0.1962 | NA |
| ABar | 11.6404 | 8506 | manual | Tintori | Xaar | 57 | 15 | 55 | 12 | 15 | NA | 12 | NA | AB | Xaa | 1.0590 | 12.7079 | NA | NA | 0.1962 | NA |
| ABal | 11.6454 | 8489 | manual | Tintori | Xaal | 57 | 15 | 55 | 12 | 15 | NA | 12 | NA | AB | Xaa | 1.0586 | 12.7037 | NA | NA | 0.1962 | NA |
| ABpr | 11.5946 | 7917 | manual | Tintori | Xapr | 57 | 15 | 55 | 12 | 15 | NA | 12 | NA | AB | Xap | 1.0569 | 12.6824 | NA | NA | 0.1962 | NA |
| ABpl | 11.7291 | 8423 | manual | Tintori | Xapl | 57 | 15 | 55 | 12 | 15 | NA | 12 | NA | AB | Xap | 1.0544 | 12.6529 | NA | NA | 0.1962 | NA |
| ABala | 10.6765 | 8628 | manual | Packer | Xaala | 74 | 17 | 74 | 17 | 17 | 15 | 12 | 15 | AB | Xaal | 1.1066 | 13.2796 | 16.5995 | 16.3782 | 0.1962 | 18.4126 |
| ABalp | 10.6765 | 8628 | manual | Packer | Xaalp | 73 | 16 | 74 | 17 | 17 | 15 | 12 | 15 | AB | Xaal | 1.1066 | 13.2796 | 16.5995 | 16.3782 | 0.1962 | 18.4126 |
| ABara | 10.6765 | 8628 | manual | Packer | Xaara | 73 | 16 | 74 | 17 | 17 | 15 | 12 | 15 | AB | Xaar | 1.1066 | 13.2796 | 16.5995 | 16.4889 | 0.1962 | 18.5370 |
| ABarp | 10.6765 | 8628 | manual | Packer | Xaarp | 73 | 16 | 73 | 17 | 17 | 15 | 12 | 15 | AB | Xaar | 1.1066 | 13.2796 | 16.5995 | 16.4889 | 0.1962 | 18.5370 |
| ABprp | 10.6136 | 7068 | script | Packer | Xaprp | 73 | 16 | 74 | 17 | 18 | 15 | 12 | 15 | AB | Xapr | 1.0976 | 13.1715 | 16.4644 | 16.3547 | 0.1962 | 18.5370 |
| ABplp | 10.6136 | 7068 | script | Packer | Xaplp | 73 | 16 | 74 | 17 | 17 | 15 | 12 | 15 | AB | Xapl | 1.0976 | 13.1715 | 16.4644 | 16.2449 | 0.1962 | 18.4126 |
| ABpra | 11.3363 | 9501 | manual | Tintori | Xapra | 73 | 16 | 74 | 17 | 17 | 15 | 12 | 15 | AB | Xapr | 1.0796 | 12.9557 | 16.1946 | 16.0866 | 0.1962 | 18.5370 |
| ABprp | 11.3363 | 9501 | manual | Tintori | Xaprp | 73 | 16 | 74 | 17 | 18 | 15 | 12 | 15 | AB | Xapr | 1.0796 | 12.9557 | 16.1946 | 16.0866 | 0.1962 | 18.5370 |
| ABpla | 11.3907 | 9393 | manual | Tintori | Xapla | 73 | 16 | 73 | 16 | 17 | 15 | 12 | 15 | AB | Xapl | 1.0764 | 12.9166 | 16.1458 | 15.9305 | 0.1962 | 18.4126 |
| ABplp | 11.3907 | 9393 | manual | Tintori | Xaplp | 73 | 16 | 74 | 17 | 17 | 15 | 12 | 15 | AB | Xapl | 1.0764 | 12.9166 | 16.1458 | 15.9305 | 0.1962 | 18.4126 |
| ABara | 11.3938 | 9792 | manual | Tintori | Xaara | 73 | 16 | 74 | 17 | 17 | 15 | 12 | 15 | AB | Xaar | 1.0787 | 12.9442 | 16.1803 | 16.0724 | 0.1962 | 18.5370 |
| ABarp | 11.3938 | 9792 | manual | Tintori | Xaarp | 73 | 16 | 73 | 17 | 17 | 15 | 12 | 15 | AB | Xaar | 1.0787 | 12.9442 | 16.1803 | 16.0724 | 0.1962 | 18.5370 |
| ABala | 11.4294 | 9602 | manual | Tintori | Xaala | 74 | 17 | 74 | 17 | 17 | 15 | 12 | 15 | AB | Xaal | 1.0759 | 12.9102 | 16.1378 | 15.9226 | 0.1962 | 18.4126 |
| ABalp | 11.4294 | 9602 | manual | Tintori | Xaalp | 73 | 16 | 74 | 17 | 17 | 15 | 12 | 15 | AB | Xaal | 1.0759 | 12.9102 | 16.1378 | 15.9226 | 0.1962 | 18.4126 |
| ABplaa | 10.6231 | 7227 | manual | Packer | Xaplaa | 96 | 23 | 100 | 26 | 24 | 17 | 16 | 16 | AB | Xapla | 1.0985 | 17.5762 | 17.5762 | 18.4550 | 0.1962 | 20.9008 |
| ABplap | 10.6231 | 7227 | manual | Packer | Xaplap | 96 | 23 | 101 | 26 | 24 | 17 | 16 | 16 | AB | Xapla | 1.0985 | 17.5762 | 17.5762 | 18.4550 | 0.1962 | 20.9008 |
| ABpraa | 10.6231 | 7227 | manual | Packer | Xapraa | 96 | 23 | 101 | 25 | 24 | 17 | 17 | 16 | AB | Xapra | 1.0985 | 18.6747 | 17.5762 | 18.7845 | 0.1962 | 21.2740 |
| ABprap | 10.6231 | 7227 | manual | Packer | Xaprap | 96 | 23 | 101 | 25 | 24 | 17 | 17 | 16 | AB | Xapra | 1.0985 | 18.6747 | 17.5762 | 18.7845 | 0.1962 | 21.2740 |
| ABplpa | 10.5976 | 6555 | script | Packer | Xaplpa | 97 | 24 | 102 | 26 | 25 | 17 | 17 | 16 | AB | Xaplp | 1.0938 | 18.5942 | 17.5004 | 19.0317 | 0.1962 | 21.6472 |
| ABplpp | 10.4747 | 5827 | script | Packer | Xaplpp | 98 | 25 | 103 | 27 | 25 | 17 | 17 | 16 | AB | Xaplp | 1.0928 | 18.5773 | 17.4845 | 19.0144 | 0.1962 | 21.6472 |
| ABprpp | 10.4747 | 5827 | script | Packer | Xaprpp | 98 | 25 | 103 | 27 | 25 | 18 | 17 | 16 | AB | Xaprp | 1.0928 | 18.5773 | 17.4845 | 19.3422 | 0.1962 | 22.0205 |
| ABprpa | 10.5976 | 6555 | script | Packer | Xaprpa | 97 | 24 | 102 | 26 | 25 | 18 | 17 | 16 | AB | Xaprp | 1.0938 | 18.5942 | 17.5004 | 19.3598 | 0.1962 | 22.0205 |
| ABarpa | 10.5369 | 7308 | script | Packer | Xaarpa | 98 | 25 | 103 | 27 | 25 | 17 | 17 | 16 | AB | Xaarp | 1.1037 | 18.7627 | 17.6590 | 18.8730 | 0.1962 | 21.2740 |
| ABarpp | 10.5369 | 7308 | script | Packer | Xaarpp | 99 | 26 | 104 | 29 | 26 | 17 | 17 | 16 | AB | Xaarp | 1.1037 | 18.7627 | 17.6590 | 18.8730 | 0.1962 | 21.2740 |
| ABarapa | 10.6008 | 6170 | script | Packer | Xaarapa | 124 | 28 | 133 | 30 | 29 | 24 | 25 | 23 | AB | Xaarap | 1.0898 | 27.2459 | 25.0662 | 26.3740 | 0.1962 | 30.1071 |
| ABarpaa | 10.2677 | 5849 | script | Packer | Xaarpaa | 122 | 24 | 131 | 27 | 26 | 25 | 27 | 25 | AB | Xaarpa | 1.1040 | 29.8074 | 27.5994 | 27.7098 | 0.1962 | 31.2268 |
| ABarpap | 10.4446 | 7065 | script | Packer | Xaarpap | 123 | 25 | 133 | 28 | 27 | 25 | 27 | 25 | AB | Xaarpa | 1.1064 | 29.8740 | 27.6611 | 27.7717 | 0.1962 | 31.2268 |
| ABarppp | 10.2791 | 7758 | script | Packer | Xaarppp | 125 | 26 | 135 | 29 | 28 | 26 | 29 | 26 | AB | Xaarpp | 1.1212 | 32.5145 | 29.1509 | 29.0388 | 0.1962 | 32.2220 |
| ABarppa | 10.2791 | 7758 | script | Packer | Xaarppa | 124 | 25 | 134 | 28 | 27 | 26 | 29 | 26 | AB | Xaarpp | 1.1212 | 32.5145 | 29.1509 | 29.0388 | 0.1962 | 32.2220 |
| ABaraaa | 10.6547 | 5987 | script | Packer | Xaaraaa | 124 | 28 | 134 | 31 | 29 | 24 | 25 | 23 | AB | Xaaraa | 1.0852 | 27.1301 | 24.9597 | 26.4789 | 0.1962 | 30.3559 |
| ABaraaa | 10.5346 | 6979 | script | Packer | Xaaraaa | 124 | 28 | 134 | 31 | 29 | 24 | 25 | 23 | AB | Xaaraa | 1.1009 | 27.5237 | 25.3218 | 26.8631 | 0.1962 | 30.3559 |
| ABaraap | 10.6547 | 5987 | script | Packer | Xaaraap | 124 | 28 | 133 | 30 | 28 | 24 | 25 | 23 | AB | Xaaraa | 1.0852 | 27.1301 | 24.9597 | 26.4789 | 0.1962 | 30.3559 |
| ABaraap | 10.5346 | 6979 | script | Packer | Xaaraap | 124 | 28 | 133 | 30 | 28 | 24 | 25 | 23 | AB | Xaaraa | 1.1009 | 27.5237 | 25.3218 | 26.8631 | 0.1962 | 30.3559 |
| ABprpap | 10.4033 | 5622 | script | Packer | Xaprpap | 124 | 27 | 133 | 29 | 28 | 25 | 26 | 24 | AB | Xaprpa | 1.0943 | 28.4507 | 26.2622 | 26.8093 | 0.1962 | 30.4803 |
| ABplpaa | 10.5346 | 5877 | script | Packer | Xaplpaa | 124 | 27 | 134 | 30 | 29 | 25 | 26 | 24 | AB | Xaplpa | 1.0902 | 28.3454 | 26.1649 | 27.1461 | 0.1962 | 30.9780 |
| ABplpap | 10.4033 | 5622 | script | Packer | Xaplpap | 123 | 26 | 132 | 29 | 28 | 25 | 26 | 24 | AB | Xaplpa | 1.0943 | 28.4507 | 26.2622 | 27.2470 | 0.1962 | 30.9780 |

| | | | | | | | | | | | | | | | | | | | | | |
|---|---|---|---|---|---|---|---|---|---|---|---|---|---|---|---|---|---|---|---|---|---|
| ABalapa | 10.3996 | 6426 | script | Packer | Xaalapa | 124 | 27 | 134 | 30 | 29 | 25 | 26 | 23 | AB | Xaalap | 1.1029 | 28.6752 | 25.3665 | 27.0208 | 0.1962 | 30.4803 |
| ABalapp | 10.3996 | 6426 | script | Packer | Xaalapp | 124 | 27 | 134 | 30 | 29 | 25 | 26 | 23 | AB | Xaalap | 1.1029 | 28.6752 | 25.3665 | 27.0208 | 0.1962 | 30.4803 |
| ABplppp | 10.5630 | 6968 | script | Packer | Xaplppp | 124 | 26 | 134 | 30 | 28 | 25 | 27 | 25 | AB | Xaplpp | 1.0994 | 29.6829 | 27.4842 | 27.8140 | 0.1962 | 31.4756 |
| ABplppa | 10.5202 | 6724 | script | Packer | Xaplppa | 126 | 28 | 135 | 31 | 30 | 25 | 27 | 25 | AB | Xaplpp | 1.0994 | 29.6833 | 27.4846 | 27.8144 | 0.1962 | 31.4756 |
| ABalaap | 10.5226 | 5157 | script | Packer | Xaalaap | 126 | 29 | 136 | 33 | 30 | 25 | 26 | 23 | AB | Xaalaa | 1.0826 | 28.1471 | 24.8994 | 26.8480 | 0.1962 | 30.8535 |
| ABalaap | 10.3591 | 6423 | script | Packer | Xaalaap | 126 | 29 | 136 | 33 | 30 | 25 | 26 | 23 | AB | Xaalaa | 1.1050 | 28.7304 | 25.4154 | 27.4044 | 0.1962 | 30.8535 |
| ABalaaa | 10.5226 | 5157 | script | Packer | Xaalaaa | 126 | 29 | 136 | 32 | 29 | 25 | 26 | 23 | AB | Xaalaa | 1.0826 | 28.1471 | 24.8994 | 26.8480 | 0.1962 | 30.8535 |
| ABalaaa | 10.3591 | 6423 | script | Packer | Xaalaaa | 126 | 29 | 136 | 32 | 29 | 25 | 26 | 23 | AB | Xaalaa | 1.1050 | 28.7304 | 25.4154 | 27.4044 | 0.1962 | 30.8535 |
| ABplapp | 10.2677 | 5825 | script | Packer | Xaplapp | 123 | 27 | 132 | 29 | 28 | 24 | 26 | 23 | AB | Xaplap | 1.1037 | 28.6967 | 25.3855 | 26.3788 | 0.1962 | 29.7339 |
| ABplapa | 10.5008 | 6187 | script | Packer | Xaplapa | 122 | 26 | 131 | 29 | 27 | 24 | 26 | 23 | AB | Xaplap | 1.0952 | 28.4748 | 25.1893 | 26.1749 | 0.1962 | 29.7339 |
| ABalpaa | 10.7045 | 7368 | script | Packer | Xaalpaa | 123 | 26 | 133 | 30 | 29 | 24 | 26 | 24 | AB | Xaalpa | 1.0955 | 28.4834 | 26.2924 | 26.6210 | 0.1962 | 30.2315 |
| ABalpaa | 10.6547 | 5987 | script | Packer | Xaalpaa | 123 | 26 | 133 | 30 | 29 | 24 | 26 | 24 | AB | Xaalpa | 1.0852 | 28.2153 | 26.0449 | 26.3704 | 0.1962 | 30.2315 |
| ABalpap | 10.7045 | 7368 | script | Packer | Xaalpap | 123 | 26 | 132 | 29 | 28 | 24 | 26 | 24 | AB | Xaalpa | 1.0955 | 28.4834 | 26.2924 | 26.6210 | 0.1962 | 30.2315 |
| ABalpap | 10.6547 | 5987 | script | Packer | Xaalpap | 123 | 26 | 132 | 29 | 28 | 24 | 26 | 24 | AB | Xaalpa | 1.0852 | 28.2153 | 26.0449 | 26.3704 | 0.1962 | 30.2315 |
| ABalppa | 10.5210 | 6523 | script | Packer | Xaalppa | 124 | 27 | 134 | 30 | 29 | 25 | 26 | 24 | AB | Xaalpp | 1.0974 | 28.5335 | 26.3387 | 27.2166 | 0.1962 | 30.8535 |
| ABalppp | 10.4436 | 7257 | script | Packer | Xaalppp | 123 | 26 | 133 | 29 | 28 | 25 | 26 | 24 | AB | Xaalpp | 1.1082 | 28.8124 | 26.5960 | 27.4826 | 0.1962 | 30.8535 |
| ABplaap | 10.5841 | 7707 | script | Packer | Xaplaap | 122 | 26 | 131 | 29 | 27 | 24 | 26 | 23 | AB | Xaplaa | 1.1045 | 28.7172 | 25.4036 | 26.6186 | 0.1962 | 29.9827 |
| ABplaaa | 10.4446 | 7065 | script | Packer | Xaplaaa | 123 | 27 | 132 | 29 | 28 | 24 | 26 | 23 | AB | Xaplaa | 1.1064 | 28.7675 | 25.4482 | 26.6653 | 0.1962 | 29.9827 |
| ABprppa | 10.5202 | 6724 | script | Packer | Xaprppa | 126 | 28 | 136 | 31 | 30 | 25 | 27 | 25 | AB | Xaprpp | 1.0994 | 29.6833 | 27.4846 | 27.1547 | 0.1962 | 30.7291 |
| ABprppp | 10.5630 | 6968 | script | Packer | Xaprppp | 125 | 27 | 134 | 30 | 29 | 25 | 27 | 25 | AB | Xaprpp | 1.0994 | 29.6829 | 27.4842 | 27.1543 | 0.1962 | 30.7291 |
| ABpraaa | 10.4436 | 7257 | script | Packer | Xapraaa | 124 | 28 | 133 | 31 | 29 | 24 | 25 | 23 | AB | Xapraa | 1.1082 | 27.7042 | 25.4879 | 27.0393 | 0.1962 | 30.3559 |
| ABpraap | 10.5841 | 7707 | script | Packer | Xapraap | 122 | 26 | 131 | 29 | 28 | 24 | 25 | 23 | AB | Xapraa | 1.1045 | 27.6126 | 25.4036 | 26.9499 | 0.1962 | 30.3559 |
| ABprpaa | 10.5346 | 5877 | script | Packer | Xaprpaa | 125 | 28 | 134 | 30 | 29 | 25 | 26 | 24 | AB | Xaprpa | 1.0902 | 28.3454 | 26.1649 | 26.7100 | 0.1962 | 30.4803 |
| ABprapp | 10.2677 | 5825 | script | Packer | Xaprapp | 123 | 27 | 132 | 30 | 28 | 24 | 25 | 23 | AB | Xaprap | 1.1037 | 27.5929 | 25.3855 | 26.5996 | 0.1962 | 29.9827 |
| ABprapa | 10.5008 | 6187 | script | Packer | Xaprapa | 123 | 27 | 132 | 29 | 27 | 24 | 25 | 23 | AB | Xaprap | 1.0952 | 27.3797 | 25.1893 | 26.3940 | 0.1962 | 29.9827 |
| ABplpaaa | 10.3636 | 7294 | script | Packer | Xaplpaaa | 157 | 33 | 172 | 36 | 34 | 29 | 30 | 27 | AB | Xaplpaa | 1.1128 | 33.3827 | 30.0445 | 31.8249 | 0.1962 | 35.5811 |
| ABplpaap | 10.4182 | 6924 | script | Packer | Xaplpaap | 156 | 32 | 171 | 35 | 34 | 29 | 30 | 27 | AB | Xaplpaa | 1.1066 | 33.1976 | 29.8778 | 31.6484 | 0.1962 | 35.5811 |
| ABplapap | 10.4787 | 7752 | script | Packer | Xaplapap | 153 | 31 | 167 | 35 | 34 | 27 | 29 | 26 | AB | Xaplapa | 1.1104 | 32.2018 | 28.8706 | 30.0921 | 0.1962 | 33.7150 |
| ABplpppp | 10.5367 | 7101 | script | Packer | Xaplpppp | 156 | 32 | 171 | 35 | 34 | 28 | 30 | 26 | AB | Xaplppp | 1.1019 | 33.0574 | 28.6498 | 31.2944 | 0.1962 | 35.3323 |
| ABplpppa | 10.3889 | 7594 | script | Packer | Xaplpppa | 157 | 33 | 172 | 36 | 35 | 28 | 30 | 26 | AB | Xaplppp | 1.1139 | 33.4175 | 28.9619 | 31.6353 | 0.1962 | 35.3323 |
| ABplpapp | 10.5640 | 6361 | script | Packer | Xaplpapp | 153 | 30 | 167 | 33 | 33 | 28 | 29 | 26 | AB | Xaplpap | 1.0936 | 31.7155 | 28.4346 | 30.0750 | 0.1962 | 34.2126 |
| ABprapap | 10.4787 | 7752 | script | Packer | Xaprapap | 155 | 32 | 169 | 36 | 34 | 27 | 29 | 27 | AB | Xaprapa | 1.1104 | 32.2018 | 29.9810 | 30.3142 | 0.1962 | 33.9638 |
| ABprapaa | 10.3475 | 7582 | script | Packer | Xaprapaa | 158 | 35 | 173 | 40 | 37 | 27 | 29 | 27 | AB | Xaprapa | 1.1160 | 32.3652 | 30.1332 | 30.4680 | 0.1962 | 33.9638 |
| ABplppaa | 10.5055 | 7309 | script | Packer | Xaplppaa | 162 | 36 | 178 | 41 | 39 | 30 | 31 | 28 | AB | Xaplppa | 1.1053 | 34.2657 | 30.9497 | 32.7182 | 0.1962 | 36.8252 |
| ABplppap | 10.3162 | 6733 | script | Packer | Xaplppap | 160 | 34 | 175 | 38 | 36 | 30 | 31 | 28 | AB | Xaplppa | 1.1103 | 34.4188 | 31.0879 | 32.8644 | 0.1962 | 36.8252 |
| ABalpapp | 10.3679 | 6829 | script | Packer | Xaalpapp | 153 | 30 | 167 | 33 | 32 | 28 | 29 | 26 | AB | Xaalpap | 1.1084 | 32.1436 | 28.8184 | 30.8135 | 0.1962 | 34.5858 |
| ABalpapa | 10.4327 | 7269 | script | Packer | Xaalpapa | 156 | 33 | 171 | 37 | 34 | 28 | 29 | 26 | AB | Xaalpap | 1.1089 | 32.1567 | 28.8301 | 30.8261 | 0.1962 | 34.5858 |
| ABarpaaa | 10.2812 | 7151 | script | Packer | Xaarpaaa | 163 | 41 | 179 | 46 | 43 | 26 | 27 | 24 | AB | Xaarpaa | 1.1160 | 30.1311 | 26.7832 | 29.1267 | 0.1962 | 32.4709 |
| ABarpaap | 10.2475 | 5847 | script | Packer | Xaarpaap | 154 | 32 | 168 | 35 | 34 | 26 | 27 | 24 | AB | Xaarpaa | 1.1050 | 29.8362 | 26.5210 | 28.8416 | 0.1962 | 32.4709 |
| ABprpapp | 10.5975 | 5860 | script | Packer | Xaprpapp | 155 | 31 | 170 | 35 | 33 | 28 | 29 | 27 | AB | Xaprpap | 1.0868 | 31.5168 | 29.3432 | 30.2126 | 0.1962 | 34.5858 |
| ABplapaa | 10.3475 | 7582 | script | Packer | Xaplapaa | 156 | 34 | 171 | 39 | 37 | 27 | 29 | 26 | AB | Xaplapa | 1.1160 | 32.3652 | 29.0171 | 30.2448 | 0.1962 | 33.7150 |
| ABalapap | 10.1627 | 7497 | script | Packer | Xaalapap | 155 | 31 | 170 | 35 | 34 | 29 | 30 | 27 | AB | Xaalapa | 1.1254 | 33.7630 | 30.3867 | 32.1874 | 0.1962 | 35.5811 |
| ABalapaa | 10.3591 | 6423 | script | Packer | Xaalapaa | 159 | 35 | 174 | 39 | 36 | 29 | 30 | 27 | AB | Xaalapa | 1.1050 | 33.1505 | 29.8354 | 31.6035 | 0.1962 | 35.5811 |

| | | | | | | | | | | | | | | | | | | | | | |
|---|---|---|---|---|---|---|---|---|---|---|---|---|---|---|---|---|---|---|---|---|---|
| ABalapaa | 10.0096 | 6057 | script | Packer | Xaalapaa | 159 | 35 | 174 | 39 | 36 | 29 | 30 | 27 | AB | Xaalapa | 1.1204 | 33.6111 | 30.2500 | 32.0426 | 0.1962 | 35.5811 |
| ABprppap | 10.3162 | 6733 | script | Packer | Xaprppap | 159 | 33 | 175 | 38 | 36 | 30 | 31 | 28 | AB | Xaprppa | 1.1103 | 34.4188 | 31.0879 | 32.7534 | 0.1962 | 36.7008 |
| ABprppaa | 10.5055 | 7309 | script | Packer | Xaprppaa | 163 | 37 | 179 | 41 | 38 | 30 | 31 | 28 | AB | Xaprppa | 1.1053 | 34.2657 | 30.9497 | 32.6077 | 0.1962 | 36.7008 |
| ABalppaa | 10.1015 | 6143 | script | Packer | Xaalppaa | 162 | 38 | 178 | 42 | 40 | 29 | 30 | 27 | AB | Xaalppa | 1.1162 | 33.4851 | 30.1366 | 32.5921 | 0.1962 | 36.3276 |
| ABalppap | 10.3317 | 7972 | script | Packer | Xaalppap | 156 | 32 | 171 | 35 | 34 | 29 | 30 | 27 | AB | Xaalppa | 1.1200 | 33.6008 | 30.2407 | 32.7048 | 0.1962 | 36.3276 |
| ABarpapp | 9.5478 | 5588 | script | Packer | Xaarpapp | 156 | 33 | 171 | 36 | 35 | 27 | 28 | 25 | AB | Xaarpap | 1.1418 | 31.9712 | 28.5457 | 31.1719 | 0.1962 | 33.9638 |
| ABarpapa | 10.0119 | 6623 | script | Packer | Xaarpapa | 155 | 32 | 170 | 36 | 34 | 27 | 28 | 25 | AB | Xaarpap | 1.1260 | 31.5273 | 28.1494 | 30.7391 | 0.1962 | 33.9638 |
| ABalpppp | 10.2923 | 6196 | script | Packer | Xaalpppp | 153 | 30 | 168 | 33 | 33 | 28 | 29 | 26 | AB | Xaalppp | 1.1063 | 32.0831 | 28.7642 | 30.7556 | 0.1962 | 34.5858 |
| ABalpppa | 10.2798 | 7342 | script | Packer | Xaalpppa | 157 | 34 | 172 | 38 | 35 | 28 | 29 | 26 | AB | Xaalppp | 1.1177 | 32.4132 | 29.0601 | 31.0719 | 0.1962 | 34.5858 |
| ABarpppp | 10.3654 | 6909 | script | Packer | Xaarpppp | 161 | 36 | 177 | 41 | 39 | 28 | 29 | 26 | AB | Xaarppp | 1.1093 | 32.1686 | 28.8408 | 30.8375 | 0.1962 | 34.5858 |
| ABplaaap | 9.5478 | 5588 | script | Packer | Xaplaaap | 156 | 33 | 171 | 38 | 35 | 28 | 29 | 27 | AB | Xaplaaa | 1.1418 | 33.1130 | 30.8294 | 31.7429 | 0.1962 | 34.5858 |
| ABplaaaa | 10.0119 | 6623 | script | Packer | Xaplaaaa | 155 | 32 | 170 | 36 | 34 | 28 | 29 | 27 | AB | Xaplaaa | 1.1260 | 32.6533 | 30.4013 | 31.3021 | 0.1962 | 34.5858 |
| ABalappa | 10.3591 | 6423 | script | Packer | Xaalappa | 162 | 38 | 177 | 42 | 39 | 29 | 30 | 27 | AB | Xaalapp | 1.1050 | 33.1505 | 29.8354 | 31.9350 | 0.1962 | 35.9543 |
| ABalappa | 9.5434 | 4235 | script | Packer | Xaalappa | 162 | 38 | 177 | 42 | 39 | 29 | 30 | 27 | AB | Xaalapp | 1.1236 | 33.7078 | 30.3370 | 32.4718 | 0.1962 | 35.9543 |
| ABalappp | 10.1627 | 7497 | script | Packer | Xaalappp | 156 | 32 | 171 | 35 | 33 | 29 | 30 | 27 | AB | Xaalapp | 1.1254 | 33.7630 | 30.3867 | 32.5250 | 0.1962 | 35.9543 |
| ABplaapa | 10.3860 | 7733 | script | Packer | Xaplaapa | 155 | 33 | 170 | 37 | 34 | 27 | 29 | 26 | AB | Xaplaap | 1.1152 | 32.3408 | 28.9952 | 29.5528 | 0.1962 | 32.9685 |
| ABplaapp | 10.5770 | 7317 | script | Packer | Xaplaapp | 152 | 30 | 166 | 34 | 32 | 27 | 29 | 26 | AB | Xaplaap | 1.1017 | 31.9484 | 28.6434 | 29.1942 | 0.1962 | 32.9685 |
| ABprpppa | 10.3889 | 7594 | script | Packer | Xaprpppa | 157 | 32 | 172 | 36 | 34 | 29 | 30 | 27 | AB | Xaprppp | 1.1139 | 33.4175 | 30.0758 | 31.8581 | 0.1962 | 35.5811 |
| ABprpppp | 10.5367 | 7101 | script | Packer | Xaprpppp | 156 | 31 | 171 | 36 | 34 | 29 | 30 | 27 | AB | Xaprppp | 1.1019 | 33.0574 | 29.7517 | 31.5147 | 0.1962 | 35.5811 |
| ABplpapa | 10.5390 | 6993 | script | Packer | Xaplpapa | 155 | 32 | 170 | 36 | 35 | 28 | 29 | 26 | AB | Xaplpap | 1.1008 | 31.9244 | 28.6219 | 30.2731 | 0.1962 | 34.2126 |
| ABalaapa | 9.9235 | 5938 | script | Packer | Xaalaapa | 164 | 38 | 180 | 43 | 39 | 30 | 33 | 29 | AB | Xaalaap | 1.1239 | 37.0900 | 32.5942 | 33.6058 | 0.1962 | 37.1984 |
| ABalaapp | 10.2176 | 6471 | script | Packer | Xaalaapp | 159 | 33 | 174 | 37 | 35 | 30 | 33 | 29 | AB | Xaalaap | 1.1131 | 36.7326 | 32.2802 | 33.2820 | 0.1962 | 37.1984 |
| ABarppap | 10.3654 | 6909 | script | Packer | Xaarppap | 161 | 37 | 177 | 41 | 39 | 27 | 28 | 25 | AB | Xaarppa | 1.1093 | 31.0594 | 27.7316 | 29.8392 | 0.1962 | 33.4661 |
| ABprpapa | 10.5390 | 6993 | script | Packer | Xaprpapa | 156 | 32 | 171 | 37 | 34 | 28 | 29 | 27 | AB | Xaprpap | 1.1008 | 31.9244 | 29.7227 | 30.6034 | 0.1962 | 34.5858 |
| ABalpaap | 10.1562 | 6202 | script | Packer | Xaalpaap | 154 | 31 | 169 | 34 | 32 | 29 | 30 | 26 | AB | Xaalpaa | 1.1138 | 33.4130 | 28.9580 | 31.7424 | 0.1962 | 35.4567 |
| ABpraaaa | 10.2798 | 7342 | script | Packer | Xapraaaa | 160 | 36 | 175 | 41 | 38 | 29 | 31 | 28 | AB | Xapraaa | 1.1177 | 34.6486 | 31.2955 | 32.6367 | 0.1962 | 36.3276 |
| ABpraaap | 10.2923 | 6196 | script | Packer | Xapraaap | 156 | 32 | 171 | 36 | 34 | 29 | 31 | 28 | AB | Xapraaa | 1.1063 | 34.2958 | 30.9768 | 32.3044 | 0.1962 | 36.3276 |
| ABarappa | 10.4327 | 7269 | script | Packer | Xaarappa | 157 | 33 | 172 | 38 | 36 | 29 | 30 | 28 | AB | Xaarapp | 1.1089 | 33.2656 | 31.0479 | 32.0458 | 0.1962 | 35.9543 |
| ABarappp | 10.3317 | 7972 | script | Packer | Xaarappp | 155 | 31 | 170 | 35 | 33 | 29 | 30 | 28 | AB | Xaarapp | 1.1200 | 33.6008 | 31.3607 | 32.3688 | 0.1962 | 35.9543 |
| ABalaaap | 10.1449 | 6677 | script | Packer | Xaalaaap | 159 | 33 | 174 | 37 | 35 | 29 | 32 | 29 | AB | Xaalaaa | 1.1191 | 35.8108 | 32.4535 | 32.7893 | 0.1962 | 36.4520 |
| ABaraaaa | 10.5291 | 7226 | script | Packer | Xaaraaaa | 162 | 38 | 178 | 43 | 39 | 29 | 31 | 28 | AB | Xaaraaa | 1.1034 | 34.2053 | 30.8951 | 32.4399 | 0.1962 | 36.5764 |
| ABaraaap | 10.1562 | 6202 | script | Packer | Xaaraaap | 156 | 32 | 171 | 35 | 33 | 29 | 31 | 28 | AB | Xaaraaa | 1.1138 | 34.5268 | 31.1855 | 32.7448 | 0.1962 | 36.5764 |
| ABalaaaa | 10.0344 | 6786 | script | Packer | Xaalaaaa | 164 | 38 | 180 | 43 | 38 | 29 | 32 | 29 | AB | Xaalaaa | 1.1263 | 36.0404 | 32.6616 | 32.9995 | 0.1962 | 36.4520 |
| ABprpaaa | 10.3636 | 7294 | script | Packer | Xaprpaaa | 157 | 32 | 172 | 37 | 34 | 29 | 30 | 28 | AB | Xaprpaa | 1.1128 | 33.3827 | 31.1572 | 31.8249 | 0.1962 | 35.5811 |
| ABprpaap | 10.4182 | 6924 | script | Packer | Xaprpaap | 156 | 31 | 171 | 35 | 34 | 29 | 30 | 28 | AB | Xaprpaa | 1.1066 | 33.1976 | 30.9844 | 31.6484 | 0.1962 | 35.5811 |
| ABplappa | 10.1405 | 7645 | script | Packer | Xaplappa | 156 | 33 | 170 | 37 | 36 | 28 | 29 | 27 | AB | Xaplapp | 1.1279 | 32.7091 | 30.4533 | 31.3556 | 0.1962 | 34.5858 |
| ABplappp | 10.2610 | 7728 | script | Packer | Xaplappp | 158 | 35 | 173 | 40 | 38 | 28 | 29 | 27 | AB | Xaplapp | 1.1219 | 32.5361 | 30.2923 | 31.1898 | 0.1962 | 34.5858 |
| ABarapap | 10.3679 | 6829 | script | Packer | Xaarapap | 154 | 30 | 168 | 34 | 33 | 29 | 30 | 28 | AB | Xaarapa | 1.1084 | 33.2520 | 31.0352 | 31.7003 | 0.1962 | 35.5811 |
| ABarapaa | 10.0007 | 5305 | script | Packer | Xaarapaa | 158 | 34 | 173 | 38 | 36 | 29 | 30 | 28 | AB | Xaarapa | 1.1123 | 33.3691 | 31.1445 | 31.8119 | 0.1962 | 35.5811 |
| ABaraapp | 10.3153 | 6516 | script | Packer | Xaaraapp | 155 | 31 | 170 | 35 | 33 | 28 | 30 | 28 | AB | Xaaraap | 1.1083 | 33.2480 | 31.0315 | 31.1423 | 0.1962 | 34.9591 |
| ABaraapa | 10.3153 | 6516 | script | Packer | Xaaraapa | 157 | 33 | 172 | 38 | 34 | 28 | 30 | 28 | AB | Xaaraap | 1.1083 | 33.2480 | 31.0315 | 31.1423 | 0.1962 | 34.9591 |
| ABpraapp | 10.5770 | 7317 | script | Packer | Xapraapp | 153 | 31 | 168 | 35 | 33 | 28 | 29 | 26 | AB | Xapraap | 1.1017 | 31.9484 | 28.6434 | 30.4061 | 0.1962 | 34.3370 |
| ABpraapa | 10.3860 | 7733 | script | Packer | Xapraapa | 156 | 34 | 171 | 38 | 35 | 28 | 29 | 26 | AB | Xapraap | 1.1152 | 32.3408 | 28.9952 | 30.7795 | 0.1962 | 34.3370 |

| | | | | | | | | | | | | | | | | | | | | | |
|---|---|---|---|---|---|---|---|---|---|---|---|---|---|---|---|---|---|---|---|---|---|
| ABprappa | 10.1405 | 7645 | script | Packer | Xaprappa | 158 | 35 | 173 | 39 | 36 | 28 | 30 | 27 | AB | Xaprapp | 1.1279 | 33.8370 | 30.4533 | 31.6940 | 0.1962 | 34.9591 |
| ABprappp | 10.2610 | 7728 | script | Packer | Xaprappp | 160 | 37 | 175 | 41 | 39 | 28 | 30 | 27 | AB | Xaprapp | 1.1219 | 33.6581 | 30.2923 | 31.5264 | 0.1962 | 34.9591 |
| ABalpaaa | 10.0007 | 5305 | script | Packer | Xaalpaaa | 161 | 38 | 176 | 42 | 38 | 29 | 30 | 26 | AB | Xaalpaa | 1.1123 | 33.3691 | 28.9199 | 31.7007 | 0.1962 | 35.4567 |
| ABalapaa | 9.9687 | 7609 | script | Packer | Xaalapaap | 201 | 42 | 224 | 48 | NA | NA | 39 | 35 | AB | Xaalapaa | 1.1373 | 44.3539 | 39.8048 | NA | 0.1962 | NA |
| ABprpapp | 10.2877 | 7173 | script | Packer | Xaprpappa | 196 | 41 | 218 | 46 | NA | NA | 35 | 31 | AB | Xaprpapp | 1.1158 | 39.0531 | 34.5899 | NA | 0.1962 | NA |
| ABalapaa | 9.7030 | 6176 | script | Packer | Xaalapaaa | 205 | 46 | 228 | 52 | NA | NA | 39 | 35 | AB | Xaalapaa | 1.1392 | 44.4290 | 39.8721 | NA | 0.1962 | NA |
| ABalapaa | 9.7028 | 6162 | script | Packer | Xaalapaaa | 205 | 46 | 228 | 52 | NA | NA | 39 | 35 | AB | Xaalapaa | 1.1391 | 44.4236 | 39.8673 | NA | 0.1962 | NA |
| ABprpapp | 10.3084 | 6723 | script | Packer | Xaprpappp | 197 | 42 | 219 | 47 | NA | NA | 35 | 31 | AB | Xaprpapp | 1.1106 | 38.8713 | 34.4289 | NA | 0.1962 | NA |
| ABarapaa | 9.8406 | 6404 | script | Packer | Xaarapaap | 197 | 39 | 218 | 44 | NA | NA | 38 | 34 | AB | Xaarapaa | 1.1336 | 43.0752 | 38.5410 | NA | 0.1962 | NA |
| ABarapaa | 9.9357 | 6635 | script | Packer | Xaarapaaa | 204 | 46 | 227 | 53 | NA | NA | 38 | 34 | AB | Xaarapaa | 1.1304 | 42.9551 | 38.4335 | NA | 0.1962 | NA |
| ABarapap | 10.1215 | 7049 | script | Packer | Xaarapapa | 194 | 40 | 216 | 46 | NA | NA | 34 | 30 | AB | Xaarapap | 1.1238 | 38.2099 | 33.7147 | NA | 0.1962 | NA |
| ABplpapa | 10.1850 | 7351 | script | Packer | Xaplpapap | 198 | 43 | 220 | 48 | NA | NA | 36 | 32 | AB | Xaplpapa | 1.1230 | 40.4266 | 35.9347 | NA | 0.1962 | NA |
| ABplpapa | 9.9841 | 7466 | script | Packer | Xaplpapaa | 200 | 45 | 223 | 51 | NA | NA | 36 | 32 | AB | Xaplpapa | 1.1352 | 40.8670 | 36.3262 | NA | 0.1962 | NA |
| ABpraapp | 10.5029 | 7248 | script | Packer | Xapraappp | 195 | 42 | 216 | 47 | NA | NA | 35 | 31 | AB | Xapraapp | 1.1050 | 38.6737 | 34.2538 | NA | 0.1962 | NA |
| ABpraapp | 10.4375 | 7368 | script | Packer | Xapraappa | 201 | 48 | 223 | 54 | NA | NA | 35 | 31 | AB | Xapraapp | 1.1094 | 38.8305 | 34.3927 | NA | 0.1962 | NA |
| ABpraaaa | 10.1691 | 7747 | script | Packer | Xapraaaap | 200 | 40 | 222 | 45 | NA | NA | 41 | 36 | AB | Xapraaaa | 1.1271 | 46.2129 | 40.5772 | NA | 0.1962 | NA |
| ABalppap | 10.1785 | 6609 | script | Packer | Xaalppapp | 195 | 39 | 216 | 44 | NA | NA | 35 | 32 | AB | Xaalppap | 1.1166 | 39.0805 | 35.7308 | NA | 0.1962 | NA |
| ABpraaaa | 10.0259 | 7639 | script | Packer | Xapraaaaa | 207 | 47 | 231 | 54 | NA | NA | 41 | 36 | AB | Xapraaaa | 1.1343 | 46.5053 | 40.8339 | NA | 0.1962 | NA |
| ABalppap | 9.9719 | 7548 | script | Packer | Xaalppapa | 197 | 41 | 219 | 46 | NA | NA | 35 | 32 | AB | Xaalppap | 1.1366 | 39.7803 | 36.3706 | NA | 0.1962 | NA |
| ABarppap | 10.1312 | 7622 | script | Packer | Xaarppapp | 211 | 50 | 235 | 56 | NA | NA | 41 | 37 | AB | Xaarppap | 1.1282 | 46.2573 | 41.7444 | NA | 0.1962 | NA |
| ABarppap | 10.1246 | 6722 | script | Packer | Xaarppapa | 210 | 49 | 234 | 55 | NA | NA | 41 | 37 | AB | Xaarppap | 1.1206 | 45.9460 | 41.4635 | NA | 0.1962 | NA |
| ABarpppa | 10.1584 | 6883 | script | Packer | Xaarpppaa | 207 | 48 | 231 | 55 | NA | NA | 38 | 34 | AB | Xaarpppa | 1.1203 | 42.5702 | 38.0891 | NA | 0.1962 | NA |
| ABarpppa | 10.1881 | 7130 | script | Packer | Xaarpppap | 202 | 43 | 225 | 49 | NA | NA | 38 | 34 | AB | Xaarpppa | 1.1209 | 42.5928 | 38.1094 | NA | 0.1962 | NA |
| ABpraapa | 10.2221 | 7499 | script | Packer | Xapraapaa | 200 | 44 | 222 | 50 | NA | NA | 38 | 34 | AB | Xapraapa | 1.1222 | 42.6427 | 38.1540 | NA | 0.1962 | NA |
| ABpraapa | 10.1436 | 7501 | script | Packer | Xapraapap | 198 | 42 | 221 | 48 | NA | NA | 38 | 34 | AB | Xapraapa | 1.1265 | 42.8080 | 38.3019 | NA | 0.1962 | NA |
| ABarpppp | 10.1246 | 6722 | script | Packer | Xaarppppa | 208 | 47 | 232 | 53 | NA | NA | 41 | 36 | AB | Xaarpppp | 1.1206 | 45.9460 | 40.3429 | NA | 0.1962 | NA |
| ABpraaap | 10.3142 | 6912 | script | Packer | Xapraaapa | 200 | 44 | 222 | 50 | NA | NA | 36 | 32 | AB | Xapraaap | 1.1120 | 40.0335 | 35.5853 | NA | 0.1962 | NA |
| ABpraaap | 10.4432 | 8121 | script | Packer | Xapraaapp | 192 | 36 | 213 | 41 | NA | NA | 36 | 32 | AB | Xapraaap | 1.1152 | 40.1465 | 35.6858 | NA | 0.1962 | NA |
| ABarppaa | 10.1584 | 6883 | script | Packer | Xaarppaaa | 209 | 49 | 233 | 56 | NA | NA | 40 | 36 | AB | Xaarppaa | 1.1203 | 44.8108 | 40.3297 | NA | 0.1962 | NA |
| ABarppaa | 10.1881 | 7130 | script | Packer | Xaarppaap | 204 | 44 | 227 | 50 | NA | NA | 40 | 36 | AB | Xaarppaa | 1.1209 | 44.8346 | 40.3511 | NA | 0.1962 | NA |
| ABalpppp | 10.4432 | 8121 | script | Packer | Xaalppppp | 189 | 36 | 210 | 40 | NA | NA | 33 | 30 | AB | Xaalpppp | 1.1152 | 36.8010 | 33.4555 | NA | 0.1962 | NA |
| ABalpppp | 10.3142 | 6912 | script | Packer | Xaalppppa | 195 | 42 | 217 | 47 | NA | NA | 33 | 30 | AB | Xaalpppp | 1.1120 | 36.6974 | 33.3612 | NA | 0.1962 | NA |
| ABplpapp | 10.5576 | 6777 | script | Packer | Xaplpappa | 191 | 38 | 212 | 43 | NA | NA | 33 | 30 | AB | Xaplpapp | 1.0979 | 36.2313 | 32.9376 | NA | 0.1962 | NA |
| ABplaaaa | 10.2849 | 6474 | script | Packer | Xaplaaaap | 199 | 44 | 221 | 50 | NA | NA | 36 | 32 | AB | Xaplaaaa | 1.1095 | 39.9418 | 35.5038 | NA | 0.1962 | NA |
| ABplaaaa | 10.1390 | 7621 | script | Packer | Xaplaaaaa | 200 | 45 | 223 | 51 | NA | NA | 36 | 32 | AB | Xaplaaaa | 1.1278 | 40.6001 | 36.0890 | NA | 0.1962 | NA |
| ABprpapa | 10.1850 | 7351 | script | Packer | Xaprpapap | 199 | 43 | 221 | 48 | NA | NA | 37 | 32 | AB | Xaprpapa | 1.1230 | 41.5495 | 35.9347 | NA | 0.1962 | NA |
| ABprpapa | 9.9841 | 7466 | script | Packer | Xaprpapaa | 202 | 46 | 225 | 52 | NA | NA | 37 | 32 | AB | Xaprpapa | 1.1352 | 42.0022 | 36.3262 | NA | 0.1962 | NA |
| ABplpppp | 10.2676 | 7429 | script | Packer | Xaplppppa | 196 | 40 | 218 | 46 | NA | NA | 35 | 32 | AB | Xaplpppp | 1.1191 | 39.1685 | 35.8112 | NA | 0.1962 | NA |
| ABplpppp | 10.3008 | 7446 | script | Packer | Xaplppppp | 199 | 43 | 221 | 49 | NA | NA | 35 | 32 | AB | Xaplpppp | 1.1174 | 39.1103 | 35.7580 | NA | 0.1962 | NA |
| ABalaaap | 9.7032 | 6529 | script | Packer | Xaalaaapa | 201 | 42 | 224 | 48 | NA | NA | 37 | 33 | AB | Xaalaaap | 1.1428 | 42.2841 | 37.7129 | NA | 0.1962 | NA |
| ABaraaaa | 9.1815 | 5370 | script | Packer | Xaaraaaap | 213 | 51 | 237 | 57 | NA | NA | 43 | 38 | AB | Xaaraaaa | 1.1617 | 49.9527 | 44.1442 | NA | 0.1962 | NA |
| ABaraaaa | 9.8941 | 5794 | script | Packer | Xaaraaaaa | 208 | 46 | 232 | 52 | NA | NA | 43 | 38 | AB | Xaaraaaa | 1.1240 | 48.3326 | 42.7125 | NA | 0.1962 | NA |
| ABalppaa | 9.6755 | 7662 | script | Packer | Xaalppaaa | 209 | 47 | 233 | 53 | NA | NA | 42 | 38 | AB | Xaalppaa | 1.1548 | 48.5028 | 43.8835 | NA | 0.1962 | NA |

| | | | | | | | | | | | | | | | | | | | | | |
|---|---|---|---|---|---|---|---|---|---|---|---|---|---|---|---|---|---|---|---|---|---|
| ABplppaa | 9.8908 | 7634 | script | Packer | Xaplppaaa | 210 | 48 | 234 | 54 | NA | NA | 41 | 36 | AB | Xaplppaa | 1.1420 | 46.8201 | 41.1103 | NA | 0.1962 | NA |
| ABplppaa | 9.9727 | 6967 | script | Packer | Xaplppaap | 207 | 45 | 230 | 50 | NA | NA | 41 | 36 | AB | Xaplppaa | 1.1314 | 46.3885 | 40.7314 | NA | 0.1962 | NA |
| ABplpppa | 10.0055 | 6590 | script | Packer | Xaplpppaa | 200 | 43 | 222 | 49 | NA | NA | 36 | 33 | AB | Xaplpppa | 1.1260 | 40.5365 | 37.1584 | NA | 0.1962 | NA |
| ABprpppp | 10.3008 | 7446 | script | Packer | Xaprppppp | 200 | 44 | 222 | 49 | NA | NA | 36 | 31 | AB | Xaprpppp | 1.1174 | 40.2277 | 34.6406 | NA | 0.1962 | NA |
| ABprpppp | 10.2676 | 7429 | script | Packer | Xaprppppa | 197 | 41 | 219 | 46 | NA | NA | 36 | 31 | AB | Xaprpppp | 1.1191 | 40.2876 | 34.6921 | NA | 0.1962 | NA |
| ABplppap | 10.2511 | 7489 | script | Packer | Xaplppapp | 206 | 46 | 230 | 53 | NA | NA | 38 | 34 | AB | Xaplppap | 1.1205 | 42.5792 | 38.0972 | NA | 0.1962 | NA |
| ABplppap | 10.2414 | 7655 | script | Packer | Xaplppapa | 203 | 43 | 226 | 49 | NA | NA | 38 | 34 | AB | Xaplppap | 1.1224 | 42.6516 | 38.1619 | NA | 0.1962 | NA |
| ABprppap | 10.2511 | 7489 | script | Packer | Xaprppapp | 206 | 47 | 230 | 53 | NA | NA | 38 | 33 | AB | Xaprppap | 1.1205 | 42.5792 | 36.9767 | NA | 0.1962 | NA |
| ABprppap | 10.2414 | 7655 | script | Packer | Xaprppapa | 202 | 43 | 225 | 49 | NA | NA | 38 | 33 | AB | Xaprppap | 1.1224 | 42.6516 | 37.0395 | NA | 0.1962 | NA |
| ABalpppa | 10.1691 | 7747 | script | Packer | Xaalpppap | 197 | 40 | 219 | 45 | NA | NA | 38 | 34 | AB | Xaalpppa | 1.1271 | 42.8315 | 38.3229 | NA | 0.1962 | NA |
| ABalpppa | 10.0259 | 7639 | script | Packer | Xaalpppaa | 203 | 46 | 226 | 52 | NA | NA | 38 | 34 | AB | Xaalpppa | 1.1343 | 43.1025 | 38.5654 | NA | 0.1962 | NA |
| ABalpapp | 10.1215 | 7049 | script | Packer | Xaalpappa | 191 | 38 | 212 | 44 | NA | NA | 33 | 30 | AB | Xaalpapp | 1.1238 | 37.0861 | 33.7147 | NA | 0.1962 | NA |
| ABalpapp | 10.4370 | 7062 | script | Packer | Xaalpappp | 191 | 38 | 212 | 43 | NA | NA | 33 | 30 | AB | Xaalpapp | 1.1068 | 36.5251 | 33.2046 | NA | 0.1962 | NA |
| ABalpaaa | 9.9357 | 6635 | script | Packer | Xaalpaaaa | 207 | 46 | 231 | 53 | NA | NA | 42 | 38 | AB | Xaalpaaa | 1.1304 | 47.4767 | 42.9551 | NA | 0.1962 | NA |
| ABalpaaa | 9.8406 | 6404 | script | Packer | Xaalpaaap | 206 | 45 | 229 | 51 | NA | NA | 42 | 38 | AB | Xaalpaaa | 1.1336 | 47.6094 | 43.0752 | NA | 0.1962 | NA |
| ABaraaap | 10.5149 | 7591 | script | Packer | Xaaraaapa | 192 | 36 | 213 | 41 | NA | NA | 35 | 32 | AB | Xaaraaap | 1.1072 | 38.7518 | 35.4303 | NA | 0.1962 | NA |
| ABalpaap | 10.1535 | 7859 | script | Packer | Xaalpaapp | 198 | 44 | 221 | 50 | NA | NA | 34 | 31 | AB | Xaalpaap | 1.1289 | 38.3831 | 34.9964 | NA | 0.1962 | NA |
| ABaraaap | 10.1535 | 7859 | script | Packer | Xaaraaapp | 202 | 46 | 224 | 52 | NA | NA | 35 | 32 | AB | Xaaraaap | 1.1289 | 39.5121 | 36.1253 | NA | 0.1962 | NA |
| ABalpaap | 10.5149 | 7591 | script | Packer | Xaalpaapa | 190 | 36 | 210 | 40 | NA | NA | 34 | 31 | AB | Xaalpaap | 1.1072 | 37.6447 | 34.3231 | NA | 0.1962 | NA |
| ABalpapa | 10.1420 | 7168 | script | Packer | Xaalpapap | 198 | 42 | 221 | 48 | NA | NA | 37 | 33 | AB | Xaalpapa | 1.1237 | 41.5786 | 37.0836 | NA | 0.1962 | NA |
| ABalpapa | 10.1100 | 7996 | script | Packer | Xaalpapaa | 199 | 43 | 221 | 49 | NA | NA | 37 | 33 | AB | Xaalpapa | 1.1324 | 41.9000 | 37.3703 | NA | 0.1962 | NA |
| ABprapap | 10.2446 | 7500 | script | Packer | Xaprapapa | 199 | 44 | 221 | 50 | NA | NA | 36 | 32 | AB | Xaprapap | 1.1209 | 40.3542 | 35.8704 | NA | 0.1962 | NA |
| ABprapap | 10.4357 | 6252 | script | Packer | Xaprapapp | 196 | 41 | 218 | 47 | NA | NA | 36 | 32 | AB | Xaprapap | 1.0993 | 39.5732 | 35.1762 | NA | 0.1962 | NA |
| ABplappa | 10.0295 | 7314 | script | Packer | Xaplappaa | 199 | 43 | 221 | 49 | NA | NA | 37 | 33 | AB | Xaplappa | 1.1313 | 41.8586 | 37.3334 | NA | 0.1962 | NA |
| ABplappa | 9.9984 | 7690 | script | Packer | Xaplappap | 199 | 43 | 221 | 49 | NA | NA | 37 | 33 | AB | Xaplappa | 1.1363 | 42.0415 | 37.4965 | NA | 0.1962 | NA |
| ABprppaa | 9.8908 | 7634 | script | Packer | Xaprppaaa | 210 | 47 | 235 | 54 | NA | NA | 41 | 37 | AB | Xaprppaa | 1.1420 | 46.8201 | 42.2523 | NA | 0.1962 | NA |
| ABplapap | 10.2446 | 7500 | script | Packer | Xaplapapa | 196 | 43 | 218 | 49 | NA | NA | 35 | 31 | AB | Xaplapap | 1.1209 | 39.2332 | 34.7494 | NA | 0.1962 | NA |
| ABprppaa | 9.9727 | 6967 | script | Packer | Xaprppaap | 206 | 43 | 230 | 50 | NA | NA | 41 | 37 | AB | Xaprppaa | 1.1314 | 46.3885 | 41.8628 | NA | 0.1962 | NA |
| ABplapap | 10.4357 | 6252 | script | Packer | Xaplapapp | 193 | 40 | 215 | 46 | NA | NA | 35 | 31 | AB | Xaplapap | 1.0993 | 38.4740 | 34.0770 | NA | 0.1962 | NA |
| ABplpaaa | 10.1335 | 7201 | script | Packer | Xaplpaaaa | 197 | 40 | 219 | 45 | NA | NA | 36 | 33 | AB | Xaplpaaa | 1.1245 | 40.4822 | 37.1087 | NA | 0.1962 | NA |
| ABplpaaa | 10.0522 | 6828 | script | Packer | Xaplpaaap | 198 | 41 | 220 | 47 | NA | NA | 36 | 33 | AB | Xaplpaaa | 1.1257 | 40.5238 | 37.1468 | NA | 0.1962 | NA |
| ABprapaa | 10.3108 | 6691 | script | Packer | Xaprapaap | 205 | 47 | 229 | 54 | NA | NA | 40 | 35 | AB | Xaprapaa | 1.1102 | 44.4071 | 38.8562 | NA | 0.1962 | NA |
| ABprapaa | 10.1578 | 7645 | script | Packer | Xaprapaaa | 202 | 44 | 224 | 50 | NA | NA | 40 | 35 | AB | Xaprapaa | 1.1269 | 45.0774 | 39.4427 | NA | 0.1962 | NA |
| ABarappa | 10.1100 | 7996 | script | Packer | Xaarappaa | 204 | 47 | 227 | 53 | NA | NA | 38 | 33 | AB | Xaarappa | 1.1324 | 43.0325 | 37.3703 | NA | 0.1962 | NA |
| ABarappa | 10.1420 | 7168 | script | Packer | Xaarappap | 200 | 43 | 222 | 48 | NA | NA | 38 | 33 | AB | Xaarappa | 1.1237 | 42.7023 | 37.0836 | NA | 0.1962 | NA |
| ABprappa | 10.0295 | 7314 | script | Packer | Xaprappaa | 202 | 44 | 225 | 50 | NA | NA | 39 | 35 | AB | Xaprappa | 1.1313 | 44.1213 | 39.5960 | NA | 0.1962 | NA |
| ABprappa | 9.9984 | 7690 | script | Packer | Xaprappap | 202 | 44 | 224 | 50 | NA | NA | 39 | 35 | AB | Xaprappa | 1.1363 | 44.3140 | 39.7690 | NA | 0.1962 | NA |
| ABaraapp | 10.1742 | 5944 | script | Packer | Xaaraappp | 195 | 40 | 217 | 46 | NA | NA | 35 | 31 | AB | Xaaraapp | 1.1101 | 38.8524 | 34.4122 | NA | 0.1962 | NA |
| ABaraapp | 10.0269 | 6748 | script | Packer | Xaaraappa | 196 | 41 | 218 | 46 | NA | NA | 35 | 31 | AB | Xaaraapp | 1.1263 | 39.4215 | 34.9162 | NA | 0.1962 | NA |
| ABalappa | 9.7762 | 5710 | script | Packer | Xaalappaa | 213 | 51 | 238 | 59 | NA | NA | 42 | 38 | AB | Xaalappa | 1.1298 | 47.4525 | 42.9332 | NA | 0.1962 | NA |
| ABalappa | 9.7028 | 6162 | script | Packer | Xaalappap | 203 | 41 | 226 | 47 | NA | NA | 42 | 38 | AB | Xaalappa | 1.1391 | 47.8408 | 43.2845 | NA | 0.1962 | NA |
| ABalappa | 9.5845 | 6270 | script | Packer | Xaalappap | 203 | 41 | 226 | 47 | NA | NA | 42 | 38 | AB | Xaalappa | 1.1472 | 48.1831 | 43.5942 | NA | 0.1962 | NA |
| ABplappp | 9.9378 | 7782 | script | Packer | Xaplapppa | 206 | 48 | 229 | 54 | NA | NA | 40 | 35 | AB | Xaplappp | 1.1405 | 45.6190 | 39.9166 | NA | 0.1962 | NA |

| | | | | | | | | | | | | | | | | | | | | | |
|---|---|---|---|---|---|---|---|---|---|---|---|---|---|---|---|---|---|---|---|---|---|
| ABplappp | 10.2066 | 6038 | script | Packer | Xaplapppp | 205 | 47 | 228 | 53 | NA | NA | 40 | 35 | AB | Xaplappp | 1.1093 | 44.3723 | 38.8258 | NA | 0.1962 | NA |
| ABarpaaa | 10.1262 | 6482 | script | Packer | Xaarpaaaa | 207 | 44 | 231 | 50 | NA | NA | 46 | 41 | AB | Xaarpaaa | 1.1182 | 51.4387 | 45.8476 | NA | 0.1962 | NA |
| ABarpaaa | 9.7788 | 6586 | script | Packer | Xaarpaaap | 212 | 49 | 236 | 56 | NA | NA | 46 | 41 | AB | Xaarpaaa | 1.1390 | 52.3917 | 46.6970 | NA | 0.1962 | NA |
| ABplapaa | 10.3108 | 6691 | script | Packer | Xaplapaap | 203 | 47 | 225 | 52 | NA | NA | 39 | 34 | AB | Xaplapaa | 1.1102 | 43.2970 | 37.7461 | NA | 0.1962 | NA |
| ABplapaa | 10.1578 | 7645 | script | Packer | Xaplapaaa | 198 | 42 | 221 | 47 | NA | NA | 39 | 34 | AB | Xaplapaa | 1.1269 | 43.9505 | 38.3158 | NA | 0.1962 | NA |
| ABarpapp | 10.0261 | 7122 | script | Packer | Xaarpappa | 204 | 48 | 227 | 55 | NA | NA | 36 | 33 | AB | Xaarpapp | 1.1298 | 40.6733 | 37.2839 | NA | 0.1962 | NA |
| ABarpapp | 10.2815 | 7329 | script | Packer | Xaarpappp | 199 | 43 | 221 | 49 | NA | NA | 36 | 33 | AB | Xaarpapp | 1.1175 | 40.2297 | 36.8772 | NA | 0.1962 | NA |
| ABprappp | 10.2066 | 6038 | script | Packer | Xaprapppp | 208 | 48 | 232 | 55 | NA | NA | 41 | 37 | AB | Xaprappp | 1.1093 | 45.4816 | 41.0444 | NA | 0.1962 | NA |
| ABprappp | 9.9378 | 7782 | script | Packer | Xaprapppa | 208 | 48 | 231 | 55 | NA | NA | 41 | 37 | AB | Xaprappp | 1.1405 | 46.7594 | 42.1975 | NA | 0.1962 | NA |
| ABaraapa | 10.1742 | 5944 | script | Packer | Xaaraapap | 197 | 40 | 218 | 45 | NA | NA | 38 | 33 | AB | Xaaraapa | 1.1101 | 42.1826 | 36.6323 | NA | 0.1962 | NA |
| ABarpaap | 10.1839 | 7253 | script | Packer | Xaarpaapa | 194 | 40 | 215 | 45 | NA | NA | 35 | 32 | AB | Xaarpaap | 1.1222 | 39.2762 | 35.9097 | NA | 0.1962 | NA |
| ABarpaap | 10.1839 | 7253 | script | Packer | Xaarpaapp | 194 | 40 | 216 | 46 | NA | NA | 35 | 32 | AB | Xaarpaap | 1.1222 | 39.2762 | 35.9097 | NA | 0.1962 | NA |
| ABarpapa | 10.2849 | 6474 | script | Packer | Xaarpapap | 199 | 44 | 222 | 50 | NA | NA | 36 | 32 | AB | Xaarpapa | 1.1095 | 39.9418 | 35.5038 | NA | 0.1962 | NA |
| ABarpapa | 10.1390 | 7621 | script | Packer | Xaarpapaa | 200 | 45 | 222 | 51 | NA | NA | 36 | 32 | AB | Xaarpapa | 1.1278 | 40.6001 | 36.0890 | NA | 0.1962 | NA |
| ABalaapa | 9.7642 | 7185 | script | Packer | Xaalaapaa | 211 | 47 | 235 | 54 | NA | NA | 43 | 38 | AB | Xaalaapa | 1.1454 | 49.2535 | 43.5264 | NA | 0.1962 | NA |
| ABalaapa | 9.7290 | 7927 | script | Packer | Xaalaapap | 210 | 46 | 234 | 52 | NA | NA | 43 | 38 | AB | Xaalaapa | 1.1538 | 49.6148 | 43.8456 | NA | 0.1962 | NA |
| ABplaapa | 10.2221 | 7499 | script | Packer | Xaplaapaa | 199 | 44 | 221 | 50 | NA | NA | 37 | 33 | AB | Xaplaapa | 1.1222 | 41.5205 | 37.0318 | NA | 0.1962 | NA |
| ABplaapa | 10.1436 | 7501 | script | Packer | Xaplaapap | 197 | 42 | 219 | 48 | NA | NA | 37 | 33 | AB | Xaplaapa | 1.1265 | 41.6814 | 37.1753 | NA | 0.1962 | NA |
| ABplpppa | 10.1291 | 7464 | script | Packer | Xaplpppap | 200 | 43 | 223 | 49 | NA | NA | 36 | 33 | AB | Xaplpppa | 1.1270 | 40.5727 | 37.1916 | NA | 0.1962 | NA |
| ABarappp | 10.1785 | 6609 | script | Packer | Xaarapppp | 196 | 41 | 218 | 46 | NA | NA | 35 | 31 | AB | Xaarappp | 1.1166 | 39.0805 | 34.6142 | NA | 0.1962 | NA |
| ABarappp | 9.9719 | 7548 | script | Packer | Xaarapppa | 198 | 43 | 220 | 49 | NA | NA | 35 | 31 | AB | Xaarappp | 1.1366 | 39.7803 | 35.2340 | NA | 0.1962 | NA |
| ABplpaap | 10.3114 | 6583 | script | Packer | Xaplpaapp | 196 | 40 | 218 | 46 | NA | NA | 35 | 32 | AB | Xaplpaap | 1.1091 | 38.8192 | 35.4919 | NA | 0.1962 | NA |
| ABalaapp | 9.9687 | 7609 | script | Packer | Xaalaappp | 199 | 40 | 222 | 46 | NA | NA | 37 | 33 | AB | Xaalaapp | 1.1373 | 42.0793 | 37.5302 | NA | 0.1962 | NA |
| ABplpaap | 10.1235 | 7276 | script | Packer | Xaplpaapa | 199 | 43 | 221 | 49 | NA | NA | 35 | 32 | AB | Xaplpaap | 1.1257 | 39.4002 | 36.0230 | NA | 0.1962 | NA |
| ABalaapp | 9.6755 | 7662 | script | Packer | Xaalaappa | 206 | 47 | 229 | 53 | NA | NA | 37 | 33 | AB | Xaalaapp | 1.1548 | 42.7287 | 38.1094 | NA | 0.1962 | NA |
| ABprpppa | 10.1291 | 7464 | script | Packer | Xaprpppap | 201 | 44 | 224 | 50 | NA | NA | 36 | 32 | AB | Xaprpppa | 1.1270 | 40.5727 | 36.0646 | NA | 0.1962 | NA |
| ABaraapa | 10.0269 | 6748 | script | Packer | Xaaraapaa | 199 | 42 | 221 | 47 | NA | NA | 38 | 33 | AB | Xaaraapa | 1.1263 | 42.8005 | 37.1689 | NA | 0.1962 | NA |
| ABalaaaa | 9.7649 | 6484 | script | Packer | Xaalaaaar | 208 | 44 | 232 | 50 | NA | NA | 43 | 38 | AB | Xaalaaaa | 1.1388 | 48.9663 | 43.2725 | NA | 0.1962 | NA |
| ABalaaaa | 9.4515 | 6131 | script | Packer | Xaalaaaal | 207 | 43 | 231 | 49 | NA | NA | 43 | 38 | AB | Xaalaaaa | 1.1538 | 49.6124 | 43.8435 | NA | 0.1962 | NA |
| ABprpaaa | 10.1335 | 7201 | script | Packer | Xaprpaaaa | 198 | 41 | 220 | 46 | NA | NA | 37 | 32 | AB | Xaprpaaa | 1.1245 | 41.6067 | 35.9842 | NA | 0.1962 | NA |
| ABalappp | 9.8564 | 6609 | script | Packer | Xaalapppp | 196 | 40 | 217 | 45 | NA | NA | 35 | 32 | AB | Xaalappp | 1.1347 | 39.7140 | 36.3100 | NA | 0.1962 | NA |
| ABprpaaa | 10.0522 | 6828 | script | Packer | Xaprpaaap | 199 | 42 | 221 | 47 | NA | NA | 37 | 32 | AB | Xaprpaaa | 1.1257 | 41.6494 | 36.0211 | NA | 0.1962 | NA |
| ABalappp | 9.9407 | 6177 | script | Packer | Xaalapppa | 196 | 40 | 218 | 45 | NA | NA | 35 | 32 | AB | Xaalappp | 1.1255 | 39.3931 | 36.0165 | NA | 0.1962 | NA |
| ABalappp | 9.8462 | 4873 | script | Packer | Xaalapppa | 196 | 40 | 218 | 45 | NA | NA | 35 | 32 | AB | Xaalappp | 1.1154 | 39.0403 | 35.6940 | NA | 0.1962 | NA |
| ABprpppa | 10.0055 | 6590 | script | Packer | Xaprpppaa | 201 | 44 | 223 | 49 | NA | NA | 36 | 32 | AB | Xaprpppa | 1.1260 | 40.5365 | 36.0324 | NA | 0.1962 | NA |
| ABalapap | 9.8835 | 5392 | script | Packer | Xaalapapa | 202 | 47 | 224 | 52 | NA | NA | 35 | 31 | AB | Xaalapap | 1.1199 | 39.1980 | 34.7182 | NA | 0.1962 | NA |
| ABalapap | 9.8462 | 4873 | script | Packer | Xaalapapa | 202 | 47 | 224 | 52 | NA | NA | 35 | 31 | AB | Xaalapap | 1.1154 | 39.0403 | 34.5786 | NA | 0.1962 | NA |
| ABalapap | 9.8564 | 6609 | script | Packer | Xaalapapp | 196 | 41 | 217 | 45 | NA | NA | 35 | 31 | AB | Xaalapap | 1.1347 | 39.7140 | 35.1753 | NA | 0.1962 | NA |
| ABarapap | 10.4370 | 7062 | script | Packer | Xaarapapp | 191 | 37 | 212 | 42 | NA | NA | 34 | 30 | AB | Xaarapap | 1.1068 | 37.6319 | 33.2046 | NA | 0.1962 | NA |
| ABarpppp | 10.1312 | 7622 | script | Packer | Xaarppppp | 209 | 48 | 233 | 55 | NA | NA | 41 | 36 | AB | Xaarpppp | 1.1282 | 46.2573 | 40.6162 | NA | 0.1962 | NA |
| ABplpapp | 10.3208 | 6802 | script | Packer | Xaplpappp | 199 | 46 | 221 | 52 | NA | NA | 33 | 30 | AB | Xaplpapp | 1.1107 | 36.6522 | 33.3202 | NA | 0.1962 | NA |
| ABplaapp | 10.5029 | 7248 | script | Packer | Xaplaappp | 192 | 40 | 213 | 45 | NA | NA | 34 | 30 | AB | Xaplaapp | 1.1050 | 37.5687 | 33.1488 | NA | 0.1962 | NA |
| ABplaapp | 10.4375 | 7368 | script | Packer | Xaplaappa | 198 | 46 | 220 | 52 | NA | NA | 34 | 30 | AB | Xaplaapp | 1.1094 | 37.7210 | 33.2832 | NA | 0.1962 | NA |

| | | | | | | | | | | | | | | | | | | | | | |
|---|---|---|---|---|---|---|---|---|---|---|---|---|---|---|---|---|---|---|---|---|---|
| ABalppaa | 9.7290 | 7927 | script | Packer | Xaalppaap | 207 | 45 | 230 | 51 | NA | NA | 42 | 38 | AB | Xaalppaa | 1.1538 | 48.4610 | 43.8456 | NA | 0.1962 | NA |
| ABprpaap | 10.3114 | 6583 | script | Packer | Xaprpaapp | 197 | 41 | 218 | 46 | NA | NA | 35 | 31 | AB | Xaprpaap | 1.1091 | 38.8192 | 34.3828 | NA | 0.1962 | NA |
| ABprpaap | 10.1235 | 7276 | script | Packer | Xaprpaapa | 200 | 44 | 222 | 50 | NA | NA | 35 | 31 | AB | Xaprpaap | 1.1257 | 39.4002 | 34.8973 | NA | 0.1962 | NA |
| ABalaaap | 9.7032 | 6529 | script | Packer | Xaalaaapp | 202 | 43 | 225 | 49 | NA | NA | 37 | 33 | AB | Xaalaaap | 1.1428 | 42.2841 | 37.7129 | NA | 0.1962 | NA |
| ABplaaap | 10.0261 | 7122 | script | Packer | Xaplaaapa | 204 | 48 | 228 | 55 | NA | NA | 38 | 33 | AB | Xaplaaap | 1.1298 | 42.9329 | 37.2839 | NA | 0.1962 | NA |
| ABplaaap | 10.2815 | 7329 | script | Packer | Xaplaaapp | 199 | 43 | 221 | 48 | NA | NA | 38 | 33 | AB | Xaplaaap | 1.1175 | 42.4647 | 36.8772 | NA | 0.1962 | NA |
| ABprpaaa | 10.2584 | 7971 | script | Packer | Xaprpaaaaa | 266 | 68 | 300 | 79 | NA | NA | 46 | 41 | AB | Xaprpaaaa | 1.1240 | 51.7047 | 46.0846 | NA | 0.1962 | NA |
| ABprpaaa | 10.5464 | 8074 | script | Packer | Xaprpaaaap | 258 | 60 | 290 | 69 | NA | NA | 46 | 41 | AB | Xaprpaaaa | 1.1094 | 51.0302 | 45.4834 | NA | 0.1962 | NA |
| ABprppaa | 10.2597 | 7738 | script | Packer | Xaprppaapp | 273 | 67 | 308 | 77 | NA | NA | 50 | 43 | AB | Xaprppaap | 1.1221 | 56.1042 | 48.2496 | NA | 0.1962 | NA |
| ABprppaa | 9.9722 | 7400 | script | Packer | Xaprppaapa | 280 | 74 | 318 | 86 | NA | NA | 50 | 43 | AB | Xaprppaap | 1.1353 | 56.7651 | 48.8180 | NA | 0.1962 | NA |
| ABarpaaa | 9.7751 | 6269 | script | Packer | Xaarpaaaaa | 287 | 80 | 327 | 95 | NA | NA | 50 | 44 | AB | Xaarpaaaa | 1.1360 | 56.7983 | 49.9825 | NA | 0.1962 | NA |
| ABarpaaa | 10.0477 | 5885 | script | Packer | Xaarpaaaap | 283 | 76 | 324 | 92 | NA | NA | 50 | 44 | AB | Xaarpaaaa | 1.1164 | 55.8199 | 49.1215 | NA | 0.1962 | NA |
| ABaraapa | 10.1732 | 6825 | script | Packer | Xaaraapapp | 277 | 80 | 312 | 92 | NA | NA | 45 | 40 | AB | Xaaraapap | 1.1189 | 50.3513 | 44.7567 | NA | 0.1962 | NA |
| ABaraapa | 10.6360 | 7816 | script | Packer | Xaaraapapa | 243 | 46 | 273 | 53 | NA | NA | 45 | 40 | AB | Xaaraapap | 1.1027 | 49.6204 | 44.1071 | NA | 0.1962 | NA |
| ABprpaaa | 10.4031 | 7181 | script | Packer | Xaprpaaapp | 258 | 59 | 290 | 67 | NA | NA | 47 | 42 | AB | Xaprpaaap | 1.1097 | 52.1542 | 46.6059 | NA | 0.1962 | NA |
| ABprpaaa | 9.9408 | 8108 | script | Packer | Xaprpaaapp | 258 | 59 | 290 | 67 | NA | NA | 47 | 42 | AB | Xaprpaaap | 1.1429 | 53.7168 | 48.0022 | NA | 0.1962 | NA |
| ABprpaaa | 9.9031 | 7085 | script | Packer | Xaprpaaapa | 280 | 81 | 316 | 94 | NA | NA | 47 | 42 | AB | Xaprpaaap | 1.1365 | 53.4141 | 47.7318 | NA | 0.1962 | NA |
| ABplapaa | 10.2533 | 7523 | script | Packer | Xaplapaaap | 265 | 67 | 298 | 76 | NA | NA | 47 | 42 | AB | Xaplapaaa | 1.1207 | 52.6714 | 47.0681 | NA | 0.1962 | NA |
| ABplapaa | 10.0829 | 7464 | script | Packer | Xaplapaaaa | 267 | 69 | 301 | 79 | NA | NA | 47 | 42 | AB | Xaplapaaa | 1.1296 | 53.0911 | 47.4431 | NA | 0.1962 | NA |
| ABprppap | 10.1155 | 7959 | script | Packer | Xaprppappp | 286 | 80 | 323 | 91 | NA | NA | 53 | 47 | AB | Xaprppapp | 1.1318 | 59.9869 | 53.1959 | NA | 0.1962 | NA |
| ABprppap | 9.8998 | 7504 | script | Packer | Xaprppappa | 273 | 67 | 308 | 77 | NA | NA | 53 | 47 | AB | Xaprppapp | 1.1403 | 60.4379 | 53.5959 | NA | 0.1962 | NA |
| ABaraaap | 10.7430 | 7422 | script | Packer | Xaaraaapap | 243 | 51 | 273 | 58 | NA | NA | 41 | 36 | AB | Xaaraaapa | 1.0940 | 44.8541 | 39.3841 | NA | 0.1962 | NA |
| ABaraaap | 10.5291 | 6946 | script | Packer | Xaaraaapaa | 259 | 67 | 291 | 76 | NA | NA | 41 | 36 | AB | Xaaraaapa | 1.1009 | 45.1386 | 39.6339 | NA | 0.1962 | NA |
| ABalpapa | 10.2310 | 7230 | script | Packer | Xaalpapapp | 266 | 68 | 300 | 78 | NA | NA | 48 | 42 | AB | Xaalpapap | 1.1194 | 53.7307 | 47.0144 | NA | 0.1962 | NA |
| ABaraapp | 10.6360 | 7816 | script | Packer | Xaaraapppa | 243 | 48 | 272 | 54 | NA | NA | 46 | 40 | AB | Xaaraappp | 1.1027 | 50.7231 | 44.1071 | NA | 0.1962 | NA |
| ABalpapa | 10.3602 | 7600 | script | Packer | Xaalpapapa | 257 | 59 | 289 | 67 | NA | NA | 48 | 42 | AB | Xaalpapap | 1.1155 | 53.5442 | 46.8512 | NA | 0.1962 | NA |
| ABaraapp | 10.1732 | 6825 | script | Packer | Xaaraapppp | 275 | 80 | 310 | 91 | NA | NA | 46 | 40 | AB | Xaaraappp | 1.1189 | 51.4703 | 44.7567 | NA | 0.1962 | NA |
| ABplaaaa | 10.2925 | 7222 | script | Packer | Xaplaaaaap | 255 | 55 | 287 | 63 | NA | NA | 51 | 45 | AB | Xaplaaaaa | 1.1160 | 56.9145 | 50.2186 | NA | 0.1962 | NA |
| ABplaaaa | 9.8444 | 6633 | script | Packer | Xaplaaaaaa | 279 | 79 | 316 | 92 | NA | NA | 51 | 45 | AB | Xaplaaaaa | 1.1356 | 57.9161 | 51.1024 | NA | 0.1962 | NA |
| ABarappp | 10.0031 | 7606 | script | Packer | Xaarappppa | 262 | 66 | 295 | 76 | NA | NA | 46 | 41 | AB | Xaarapppp | 1.1353 | 52.2235 | 46.5471 | NA | 0.1962 | NA |
| ABarappp | 10.1720 | 7073 | script | Packer | Xaarappppp | 254 | 58 | 286 | 67 | NA | NA | 46 | 41 | AB | Xaarapppp | 1.1212 | 51.5772 | 45.9710 | NA | 0.1962 | NA |
| ABarappp | 10.1083 | 7285 | script | Packer | Xaarappppp | 254 | 58 | 286 | 67 | NA | NA | 46 | 41 | AB | Xaarapppp | 1.1266 | 51.8257 | 46.1924 | NA | 0.1962 | NA |
| ABpraapa | 10.2430 | 7525 | script | Packer | Xapraapapa | 273 | 75 | 307 | 85 | NA | NA | 48 | 42 | AB | Xapraapap | 1.1212 | 53.8198 | 47.0923 | NA | 0.1962 | NA |
| ABalappa | 10.1873 | 7499 | script | Packer | Xaalappaap | 300 | 87 | 338 | 98 | NA | NA | 59 | 51 | AB | Xaalappaa | 1.1241 | 66.3214 | 57.3287 | NA | 0.1962 | NA |
| ABplpppa | 10.2193 | 7492 | script | Packer | Xaplpppaap | 266 | 66 | 299 | 75 | NA | NA | 49 | 43 | AB | Xaplpppaa | 1.1223 | 54.9914 | 48.2578 | NA | 0.1962 | NA |
| ABplpppa | 10.2821 | 7840 | script | Packer | Xaplpppaaa | 269 | 69 | 303 | 79 | NA | NA | 49 | 43 | AB | Xaplpppaa | 1.1217 | 54.9623 | 48.2323 | NA | 0.1962 | NA |
| ABalpaaa | 10.1326 | 7216 | script | Packer | Xaalpaaaaa | 272 | 65 | 306 | 74 | NA | NA | 53 | 46 | AB | Xaalpaaaa | 1.1247 | 59.6084 | 51.7356 | NA | 0.1962 | NA |
| ABalpaaa | 10.0594 | 6919 | script | Packer | Xaalpaaaap | 280 | 73 | 317 | 84 | NA | NA | 53 | 46 | AB | Xaalpaaaa | 1.1261 | 59.6833 | 51.8006 | NA | 0.1962 | NA |
| ABalappp | 9.8919 | 7407 | script | Packer | Xaalappppa | 246 | 50 | 277 | 58 | NA | NA | 45 | 40 | AB | Xaalapppp | 1.1400 | 51.2983 | 45.5985 | NA | 0.1962 | NA |
| ABalappp | 10.1720 | 7073 | script | Packer | Xaalappppa | 246 | 50 | 277 | 58 | NA | NA | 45 | 40 | AB | Xaalapppp | 1.1212 | 50.4560 | 44.8498 | NA | 0.1962 | NA |
| ABalappp | 9.9903 | 7078 | script | Packer | Xaalappppp | 258 | 62 | 290 | 71 | NA | NA | 45 | 40 | AB | Xaalapppp | 1.1314 | 50.9146 | 45.2575 | NA | 0.1962 | NA |
| ABalappp | 10.1833 | 7541 | script | Packer | Xaalapppaa | 261 | 65 | 294 | 75 | NA | NA | 45 | 40 | AB | Xaalapppa | 1.1247 | 50.6098 | 44.9865 | NA | 0.1962 | NA |
| ABaraaaa | 9.8454 | 7011 | script | Packer | Xaaraaaaaa | 279 | 71 | 314 | 81 | NA | NA | 52 | 46 | AB | Xaaraaaaa | 1.1391 | 59.2343 | 52.3996 | NA | 0.1962 | NA |

| | | | | | | | | | | | | | | | | | | | | | |
|---|---|---|---|---|---|---|---|---|---|---|---|---|---|---|---|---|---|---|---|---|---|
| ABarappp | 10.1590 | 7128 | script | Packer | Xaarapppap | 265 | 67 | 299 | 77 | NA | NA | 49 | 43 | AB | Xaarapppa | 1.1225 | 55.0002 | 48.2655 | NA | 0.1962 | NA |
| ABarappp | 9.8623 | 7619 | script | Packer | Xaarapppaa | 269 | 71 | 304 | 82 | NA | NA | 49 | 43 | AB | Xaarapppa | 1.1435 | 56.0305 | 49.1696 | NA | 0.1962 | NA |
| ABalappp | 10.0949 | 6266 | script | Packer | Xaalapppap | 251 | 55 | 283 | 64 | NA | NA | 45 | 40 | AB | Xaalapppa | 1.1178 | 50.3009 | 44.7119 | NA | 0.1962 | NA |
| ABalappp | 10.4031 | 7181 | script | Packer | Xaalapppap | 251 | 55 | 283 | 64 | NA | NA | 45 | 40 | AB | Xaalapppa | 1.1097 | 49.9349 | 44.3866 | NA | 0.1962 | NA |
| ABalpppa | 10.3724 | 8121 | script | Packer | Xaalpppapa | 263 | 66 | 297 | 77 | NA | NA | 45 | 40 | AB | Xaalpppap | 1.1190 | 50.3542 | 44.7593 | NA | 0.1962 | NA |
| ABalpppa | 10.2715 | 7252 | script | Packer | Xaalpppapp | 253 | 56 | 286 | 65 | NA | NA | 45 | 40 | AB | Xaalpppap | 1.1174 | 50.2816 | 44.6948 | NA | 0.1962 | NA |
| ABalappa | 9.5992 | 7288 | script | Packer | Xaalappapp | 261 | 58 | 294 | 66 | NA | NA | 47 | 41 | AB | Xaalappap | 1.1562 | 54.3395 | 47.4026 | NA | 0.1962 | NA |
| ABalappa | 9.4037 | 7827 | script | Packer | Xaalappapa | 291 | 88 | 329 | 101 | NA | NA | 47 | 41 | AB | Xaalappap | 1.1728 | 55.1211 | 48.0844 | NA | 0.1962 | NA |
| ABarappa | 10.1136 | 6952 | script | Packer | Xaarappaaa | 273 | 69 | 308 | 79 | NA | NA | 53 | 47 | AB | Xaarappaa | 1.1234 | 59.5393 | 52.7990 | NA | 0.1962 | NA |
| ABplpapa | 10.3308 | 7583 | script | Packer | Xaplpapapp | 260 | 62 | 293 | 71 | NA | NA | 48 | 43 | AB | Xaplpapap | 1.1170 | 53.6137 | 48.0290 | NA | 0.1962 | NA |
| ABarappa | 10.1397 | 6601 | script | Packer | Xaarappaap | 279 | 75 | 315 | 86 | NA | NA | 53 | 47 | AB | Xaarappaa | 1.1186 | 59.2881 | 52.5763 | NA | 0.1962 | NA |
| ABplpapa | 10.3672 | 7811 | script | Packer | Xaplpapapa | 258 | 60 | 291 | 69 | NA | NA | 48 | 43 | AB | Xaplpapap | 1.1168 | 53.6083 | 48.0241 | NA | 0.1962 | NA |
| ABalaaap | 9.8651 | 7635 | script | Packer | Xaalaaapal | 264 | 63 | 298 | 72 | NA | NA | 48 | 42 | AB | Xaalaaapa | 1.1434 | 54.8856 | 48.0249 | NA | 0.1962 | NA |
| ABalaaap | 9.2827 | 7628 | script | Packer | Xaalaaapar | 276 | 75 | 312 | 87 | NA | NA | 48 | 42 | AB | Xaalaaapa | 1.1787 | 56.5782 | 49.5059 | NA | 0.1962 | NA |
| ABprpapa | 10.3308 | 7583 | script | Packer | Xaprpapapp | 260 | 61 | 292 | 69 | NA | NA | 48 | 43 | AB | Xaprpapap | 1.1170 | 53.6137 | 48.0290 | NA | 0.1962 | NA |
| ABprpapa | 10.3672 | 7811 | script | Packer | Xaprpapapa | 257 | 58 | 290 | 67 | NA | NA | 48 | 43 | AB | Xaprpapap | 1.1168 | 53.6083 | 48.0241 | NA | 0.1962 | NA |
| ABarpaaa | 10.0477 | 5885 | script | Packer | Xaarpaaapa | 284 | 72 | 324 | 87 | NA | NA | 56 | 49 | AB | Xaarpaaap | 1.1164 | 62.5182 | 54.7035 | NA | 0.1962 | NA |
| ABarappa | 10.3602 | 7600 | script | Packer | Xaarappapa | 261 | 61 | 294 | 71 | NA | NA | 48 | 43 | AB | Xaarappap | 1.1155 | 53.5442 | 47.9667 | NA | 0.1962 | NA |
| ABarappa | 10.2310 | 7230 | script | Packer | Xaarappapp | 267 | 67 | 300 | 77 | NA | NA | 48 | 43 | AB | Xaarappap | 1.1194 | 53.7307 | 48.1338 | NA | 0.1962 | NA |
| ABalpppa | 9.9506 | 7297 | script | Packer | Xaalpppaap | 281 | 78 | 318 | 90 | NA | NA | 52 | 46 | AB | Xaalpppaa | 1.1356 | 59.0534 | 52.2396 | NA | 0.1962 | NA |
| ABalpppa | 9.6221 | 7446 | script | Packer | Xaalpppaaa | 277 | 74 | 313 | 85 | NA | NA | 52 | 46 | AB | Xaalpppaa | 1.1562 | 60.1212 | 53.1842 | NA | 0.1962 | NA |
| ABplpapp | 10.2863 | 5815 | script | Packer | Xaplpapppp | 269 | 70 | 304 | 81 | NA | NA | 52 | 46 | AB | Xaplpappp | 1.1026 | 57.3357 | 50.7200 | NA | 0.1962 | NA |
| ABplpapp | 10.3017 | 6165 | script | Packer | Xaplpapppa | 266 | 67 | 299 | 77 | NA | NA | 52 | 46 | AB | Xaplpappp | 1.1055 | 57.4857 | 50.8528 | NA | 0.1962 | NA |
| ABprppap | 10.2344 | 8384 | script | Packer | Xaprppapaa | 262 | 60 | 295 | 68 | NA | NA | 49 | 43 | AB | Xaprppapa | 1.1285 | 55.2960 | 48.5251 | NA | 0.1962 | NA |
| ABprppap | 10.1417 | 7431 | script | Packer | Xaprppapap | 281 | 79 | 317 | 91 | NA | NA | 49 | 43 | AB | Xaprppapa | 1.1260 | 55.1760 | 48.4197 | NA | 0.1962 | NA |
| ABplpapa | 10.2182 | 8062 | script | Packer | Xaplpapaaa | 268 | 68 | 302 | 77 | NA | NA | 51 | 45 | AB | Xaplpapaa | 1.1269 | 57.4736 | 50.7120 | NA | 0.1962 | NA |
| ABaraaap | 10.1873 | 5633 | script | Packer | Xaaraaappp | 268 | 66 | 302 | 76 | NA | NA | 52 | 46 | AB | Xaaraaapp | 1.1059 | 57.5079 | 50.8724 | NA | 0.1962 | NA |
| ABplpapa | 10.1385 | 7653 | script | Packer | Xaplpapaap | 269 | 69 | 304 | 79 | NA | NA | 51 | 45 | AB | Xaplpapaa | 1.1281 | 57.5319 | 50.7634 | NA | 0.1962 | NA |
| ABaraaap | 10.0287 | 6762 | script | Packer | Xaaraaappa | 278 | 76 | 314 | 88 | NA | NA | 52 | 46 | AB | Xaaraaapp | 1.1264 | 58.5706 | 51.8125 | NA | 0.1962 | NA |
| ABalpapa | 10.3774 | 6767 | script | Packer | Xaalpapaaa | 257 | 58 | 289 | 66 | NA | NA | 49 | 43 | AB | Xaalpapaa | 1.1073 | 54.2586 | 47.6147 | NA | 0.1962 | NA |
| ABalpapa | 10.1397 | 6601 | script | Packer | Xaalpapaap | 274 | 75 | 309 | 86 | NA | NA | 49 | 43 | AB | Xaalpapaa | 1.1186 | 54.8135 | 48.1017 | NA | 0.1962 | NA |
| ABaraapp | 10.3066 | 7255 | script | Packer | Xaaraappap | 251 | 55 | 283 | 63 | NA | NA | 46 | 41 | AB | Xaaraappa | 1.1155 | 51.3128 | 45.7353 | NA | 0.1962 | NA |
| ABaraapp | 9.9629 | 5567 | script | Packer | Xaaraappaa | 268 | 72 | 302 | 83 | NA | NA | 46 | 41 | AB | Xaaraappa | 1.1175 | 51.4070 | 45.8193 | NA | 0.1962 | NA |
| ABalpaap | 10.7430 | 7422 | script | Packer | Xaalpaapap | 233 | 43 | 262 | 50 | NA | NA | 40 | 36 | AB | Xaalpaapa | 1.0940 | 43.7601 | 39.3841 | NA | 0.1962 | NA |
| ABarpapa | 10.2925 | 7222 | script | Packer | Xaarpapaap | 255 | 55 | 286 | 62 | NA | NA | 51 | 45 | AB | Xaarpapaa | 1.1160 | 56.9145 | 50.2186 | NA | 0.1962 | NA |
| ABarpapa | 9.8444 | 6633 | script | Packer | Xaarpapaaa | 278 | 78 | 315 | 91 | NA | NA | 51 | 45 | AB | Xaarpapaa | 1.1356 | 57.9161 | 51.1024 | NA | 0.1962 | NA |
| ABpraapa | 10.5447 | 8138 | script | Packer | Xapraapaap | 261 | 61 | 294 | 70 | NA | NA | 50 | 44 | AB | Xapraapaa | 1.1099 | 55.4963 | 48.8368 | NA | 0.1962 | NA |
| ABprpapp | 10.3669 | 6531 | script | Packer | Xaprpappap | 256 | 60 | 288 | 69 | NA | NA | 46 | 41 | AB | Xaprpappa | 1.1056 | 50.8597 | 45.3315 | NA | 0.1962 | NA |
| ABprpapp | 10.4442 | 6449 | script | Packer | Xaprpappaa | 255 | 59 | 286 | 67 | NA | NA | 46 | 41 | AB | Xaprpappa | 1.1008 | 50.6347 | 45.1309 | NA | 0.1962 | NA |
| ABpraapa | 10.3682 | 7879 | script | Packer | Xapraapaaa | 271 | 71 | 306 | 82 | NA | NA | 50 | 44 | AB | Xapraapaa | 1.1173 | 55.8663 | 49.1623 | NA | 0.1962 | NA |
| ABplappp | 9.8431 | 7656 | script | Packer | Xaplapppaa | 286 | 80 | 326 | 95 | NA | NA | 54 | 48 | AB | Xaplapppa | 1.1449 | 61.8247 | 54.9553 | NA | 0.1962 | NA |
| ABplappp | 10.1486 | 7050 | script | Packer | Xaplapppap | 266 | 60 | 300 | 69 | NA | NA | 54 | 48 | AB | Xaplapppa | 1.1223 | 60.6057 | 53.8717 | NA | 0.1962 | NA |
| ABaraaaa | 9.8112 | 6096 | script | Packer | Xaaraaaapa | 292 | 79 | 329 | 91 | NA | NA | 57 | 51 | AB | Xaaraaaap | 1.1321 | 64.5274 | 57.7351 | NA | 0.1962 | NA |

| | | | | | | | | | | | | | | | | | | | | | |
|---|---|---|---|---|---|---|---|---|---|---|---|---|---|---|---|---|---|---|---|---|---|
| ABplaapa | 10.5447 | 8138 | script | Packer | Xaplaapaap | 261 | 62 | 294 | 71 | NA | NA | 50 | 44 | AB | Xaplaapaa | 1.1099 | 55.4963 | 48.8368 | NA | 0.1962 | NA |
| ABplaapa | 10.3682 | 7879 | script | Packer | Xaplaapaaa | 271 | 72 | 306 | 83 | NA | NA | 50 | 44 | AB | Xaplaapaa | 1.1173 | 55.8663 | 49.1623 | NA | 0.1962 | NA |
| ABprappp | 10.1486 | 7050 | script | Packer | Xaprapppap | 270 | 62 | 305 | 72 | NA | NA | 55 | 48 | AB | Xaprapppa | 1.1223 | 61.7280 | 53.8717 | NA | 0.1962 | NA |
| ABprappp | 9.8431 | 7656 | script | Packer | Xaprapppaa | 291 | 83 | 332 | 99 | NA | NA | 55 | 48 | AB | Xaprapppa | 1.1449 | 62.9696 | 54.9553 | NA | 0.1962 | NA |
| ABplaapa | 10.2430 | 7525 | script | Packer | Xaplaapapa | 272 | 75 | 307 | 86 | NA | NA | 48 | 42 | AB | Xaplaapap | 1.1212 | 53.8198 | 47.0923 | NA | 0.1962 | NA |
| ABalpaaa | 9.9706 | 5741 | script | Packer | Xaalpaaapp | 277 | 71 | 312 | 82 | NA | NA | 51 | 45 | AB | Xaalpaaap | 1.1191 | 57.0742 | 50.3596 | NA | 0.1962 | NA |
| ABalpaaa | 10.0620 | 6464 | script | Packer | Xaalpaaapa | 277 | 71 | 313 | 83 | NA | NA | 51 | 45 | AB | Xaalpaaap | 1.1216 | 57.2023 | 50.4726 | NA | 0.1962 | NA |
| ABpraaaa | 10.3724 | 8121 | script | Packer | Xapraaaapa | 268 | 68 | 302 | 78 | NA | NA | 45 | 40 | AB | Xapraaaap | 1.1190 | 50.3542 | 44.7593 | NA | 0.1962 | NA |
| ABplaapa | 10.2876 | 7327 | script | Packer | Xaplaapapp | 253 | 56 | 284 | 63 | NA | NA | 48 | 42 | AB | Xaplaapap | 1.1171 | 53.6228 | 46.9200 | NA | 0.1962 | NA |
| ABpraaaa | 10.2715 | 7252 | script | Packer | Xapraaaapp | 259 | 59 | 292 | 68 | NA | NA | 45 | 40 | AB | Xapraaaap | 1.1174 | 50.2816 | 44.6948 | NA | 0.1962 | NA |
| ABalpaap | 10.0287 | 6762 | script | Packer | Xaalpaappa | 273 | 75 | 308 | 86 | NA | NA | 50 | 44 | AB | Xaalpaapp | 1.1264 | 56.3179 | 49.5597 | NA | 0.1962 | NA |
| ABplppaa | 10.2790 | 7719 | script | Packer | Xaplppaaaa | 263 | 53 | 296 | 60 | NA | NA | 54 | 48 | AB | Xaplppaaa | 1.1209 | 60.5275 | 53.8022 | NA | 0.1962 | NA |
| ABalpaap | 10.2220 | 6433 | script | Packer | Xaalpaappp | 263 | 65 | 296 | 74 | NA | NA | 50 | 44 | AB | Xaalpaapp | 1.1125 | 55.6248 | 48.9498 | NA | 0.1962 | NA |
| ABprpapp | 10.0130 | 6597 | script | Packer | Xaprpapppa | 260 | 63 | 293 | 73 | NA | NA | 47 | 42 | AB | Xaprpappp | 1.1257 | 52.9060 | 47.2777 | NA | 0.1962 | NA |
| ABprpapp | 10.3244 | 5951 | script | Packer | Xaprpapppp | 259 | 62 | 291 | 71 | NA | NA | 47 | 42 | AB | Xaprpappp | 1.1020 | 51.7959 | 46.2857 | NA | 0.1962 | NA |
| ABalaaaa | 9.7790 | 7450 | script | Packer | Xaalaaaarr | 278 | 70 | 313 | 80 | NA | NA | 50 | 44 | AB | Xaalaaaar | 1.1469 | 57.3447 | 50.4633 | NA | 0.1962 | NA |
| ABalaaaa | 9.7790 | 7450 | script | Packer | Xaalaaaarl | 278 | 70 | 313 | 80 | NA | NA | 50 | 44 | AB | Xaalaaaar | 1.1469 | 57.3447 | 50.4633 | NA | 0.1962 | NA |
| ABalaaaa | 9.4747 | 6661 | script | Packer | Xaalaaaala | 289 | 82 | 329 | 97 | NA | NA | 49 | 43 | AB | Xaalaaaal | 1.1578 | 56.7337 | 49.7867 | NA | 0.1962 | NA |
| ABalaaaa | 9.8061 | 7232 | script | Packer | Xaalaaaalp | 287 | 80 | 324 | 92 | NA | NA | 49 | 43 | AB | Xaalaaaal | 1.1434 | 56.0266 | 49.1662 | NA | 0.1962 | NA |
| ABalppap | 10.1590 | 7128 | script | Packer | Xaalppapap | 264 | 67 | 298 | 77 | NA | NA | 46 | 41 | AB | Xaalppapa | 1.1225 | 51.6328 | 46.0206 | NA | 0.1962 | NA |
| ABpraaap | 10.1498 | 7775 | script | Packer | Xapraaapaa | 273 | 73 | 308 | 84 | NA | NA | 50 | 44 | AB | Xapraaapa | 1.1284 | 56.4223 | 49.6516 | NA | 0.1962 | NA |
| ABplpaaa | 10.2584 | 7971 | script | Packer | Xaplpaaaaa | 264 | 67 | 298 | 77 | NA | NA | 45 | 40 | AB | Xaplpaaaa | 1.1240 | 50.5807 | 44.9606 | NA | 0.1962 | NA |
| ABalaaap | 9.8651 | 7635 | script | Packer | Xaalaaappr | 266 | 64 | 300 | 74 | NA | NA | 49 | 43 | AB | Xaalaaapp | 1.1434 | 56.0290 | 49.1683 | NA | 0.1962 | NA |
| ABalaaap | 9.2827 | 7628 | script | Packer | Xaalaaappl | 276 | 74 | 312 | 86 | NA | NA | 49 | 43 | AB | Xaalaaapp | 1.1787 | 57.7569 | 50.6846 | NA | 0.1962 | NA |
| ABpraaap | 10.1227 | 7505 | script | Packer | Xapraaapap | 264 | 64 | 298 | 74 | NA | NA | 50 | 44 | AB | Xapraaapa | 1.1277 | 56.3861 | 49.6198 | NA | 0.1962 | NA |
| ABplpaaa | 10.5464 | 8074 | script | Packer | Xaplpaaaap | 257 | 60 | 289 | 69 | NA | NA | 45 | 40 | AB | Xaplpaaaa | 1.1094 | 49.9208 | 44.3741 | NA | 0.1962 | NA |
| ABpraaaa | 9.9506 | 7297 | script | Packer | Xapraaaaap | 286 | 79 | 327 | 94 | NA | NA | 54 | 47 | AB | Xapraaaaa | 1.1356 | 61.3247 | 53.3752 | NA | 0.1962 | NA |
| ABpraaaa | 9.6221 | 7446 | script | Packer | Xapraaaaaa | 279 | 72 | 316 | 83 | NA | NA | 54 | 47 | AB | Xapraaaaa | 1.1562 | 62.4336 | 54.3403 | NA | 0.1962 | NA |
| ABplpaap | 10.2240 | 7840 | script | Packer | Xaplpaappp | 256 | 60 | 288 | 69 | NA | NA | 46 | 40 | AB | Xaplpaapp | 1.1249 | 51.7438 | 44.9946 | NA | 0.1962 | NA |
| ABprpppp | 10.4825 | 7404 | script | Packer | Xaprppppa | 274 | 74 | 309 | 86 | NA | NA | 49 | 44 | AB | Xaprppppp | 1.1074 | 54.2606 | 48.7238 | NA | 0.1962 | NA |
| ABprpppp | 10.1514 | 6959 | script | Packer | Xaprppppp | 259 | 59 | 292 | 69 | NA | NA | 49 | 44 | AB | Xaprppppp | 1.1214 | 54.9462 | 49.3395 | NA | 0.1962 | NA |
| ABalapaa | 10.1274 | 7353 | script | Packer | Xaalapaapp | 258 | 57 | 290 | 65 | NA | NA | 48 | 42 | AB | Xaalapaap | 1.1262 | 54.0561 | 47.2991 | NA | 0.1962 | NA |
| ABalapaa | 10.1720 | 7073 | script | Packer | Xaalapaapp | 258 | 57 | 290 | 65 | NA | NA | 48 | 42 | AB | Xaalapaap | 1.1212 | 53.8197 | 47.0923 | NA | 0.1962 | NA |
| ABalapaa | 9.8916 | 7606 | script | Packer | Xaalapaapa | 282 | 81 | 319 | 94 | NA | NA | 48 | 42 | AB | Xaalapaap | 1.1417 | 54.8003 | 47.9502 | NA | 0.1962 | NA |
| ABalapap | 10.0949 | 6266 | script | Packer | Xaalapapap | 257 | 55 | 289 | 63 | NA | NA | 52 | 47 | AB | Xaalapapa | 1.1178 | 58.1254 | 52.5365 | NA | 0.1962 | NA |
| ABalapap | 10.4031 | 7181 | script | Packer | Xaalapapap | 257 | 55 | 289 | 63 | NA | NA | 52 | 47 | AB | Xaalapapa | 1.1097 | 57.7026 | 52.1542 | NA | 0.1962 | NA |
| ABprapap | 10.2151 | 8925 | script | Packer | Xaprapappp | 275 | 79 | 310 | 91 | NA | NA | 47 | 41 | AB | Xaprapapp | 1.1335 | 53.2725 | 46.4718 | NA | 0.1962 | NA |
| ABplpaaa | 9.9031 | 7085 | script | Packer | Xaplpaaapa | 280 | 82 | 317 | 95 | NA | NA | 47 | 41 | AB | Xaplpaaap | 1.1365 | 53.4141 | 46.5953 | NA | 0.1962 | NA |
| ABpraaap | 10.4289 | 7798 | script | Packer | Xapraaappp | 238 | 46 | 267 | 52 | NA | NA | 41 | 36 | AB | Xapraaapp | 1.1134 | 45.6504 | 40.0833 | NA | 0.1962 | NA |
| ABplpaaa | 10.4031 | 7181 | script | Packer | Xaplpaaapp | 256 | 58 | 288 | 66 | NA | NA | 47 | 41 | AB | Xaplpaaap | 1.1097 | 52.1542 | 45.4963 | NA | 0.1962 | NA |
| ABplpaaa | 9.9408 | 8108 | script | Packer | Xaplpaaapp | 256 | 58 | 288 | 66 | NA | NA | 47 | 41 | AB | Xaplpaaap | 1.1429 | 53.7168 | 46.8593 | NA | 0.1962 | NA |
| ABpraaap | 10.0810 | 7416 | script | Packer | Xapraaappa | 261 | 69 | 294 | 80 | NA | NA | 41 | 36 | AB | Xapraaapp | 1.1293 | 46.3011 | 40.6546 | NA | 0.1962 | NA |
| ABprpppp | 10.5624 | 8080 | script | Packer | Xaprppppap | 257 | 60 | 289 | 69 | NA | NA | 46 | 41 | AB | Xaprppppa | 1.1086 | 50.9937 | 45.4509 | NA | 0.1962 | NA |

| | | | | | | | | | | | | | | | | | | | | | |
|---|---|---|---|---|---|---|---|---|---|---|---|---|---|---|---|---|---|---|---|---|---|
| ABprpppp | 10.5519 | 8433 | script | Packer | Xaprppppaa | 260 | 63 | 293 | 72 | NA | NA | 46 | 41 | AB | Xaprppppa | 1.1117 | 51.1401 | 45.5814 | NA | 0.1962 | NA |
| ABplpaap | 10.3265 | 7420 | script | Packer | Xaplpaapap | 265 | 66 | 299 | 76 | NA | NA | 49 | 43 | AB | Xaplpaapa | 1.1158 | 54.6756 | 47.9806 | NA | 0.1962 | NA |
| ABplpaap | 10.0720 | 7996 | script | Packer | Xaplpaapaa | 265 | 66 | 298 | 75 | NA | NA | 49 | 43 | AB | Xaplpaapa | 1.1346 | 55.5937 | 48.7863 | NA | 0.1962 | NA |
| ABalaapa | 9.8061 | 7232 | script | Packer | Xaalaapaap | 287 | 76 | 325 | 88 | NA | NA | 54 | 47 | AB | Xaalaapaa | 1.1434 | 61.7436 | 53.7398 | NA | 0.1962 | NA |
| ABalaapa | 9.4747 | 6661 | script | Packer | Xaalaapaaa | 294 | 83 | 334 | 98 | NA | NA | 54 | 47 | AB | Xaalaapaa | 1.1578 | 62.5228 | 54.4180 | NA | 0.1962 | NA |
| ABalppap | 10.1720 | 7073 | script | Packer | Xaalppappp | 254 | 59 | 286 | 67 | NA | NA | 44 | 39 | AB | Xaalppapp | 1.1212 | 49.3347 | 43.7285 | NA | 0.1962 | NA |
| ABalppap | 10.1083 | 7285 | script | Packer | Xaalppappp | 254 | 59 | 286 | 67 | NA | NA | 44 | 39 | AB | Xaalppapp | 1.1266 | 49.5724 | 43.9391 | NA | 0.1962 | NA |
| ABalppap | 10.0031 | 7606 | script | Packer | Xaalppappa | 262 | 67 | 295 | 77 | NA | NA | 44 | 39 | AB | Xaalppapp | 1.1353 | 49.9529 | 44.2765 | NA | 0.1962 | NA |
| ABalpppp | 10.0810 | 7416 | script | Packer | Xaalppppa | 253 | 64 | 284 | 73 | NA | NA | 40 | 36 | AB | Xaalppppp | 1.1293 | 45.1718 | 40.6546 | NA | 0.1962 | NA |
| ABalpppp | 10.4289 | 7798 | script | Packer | Xaalpppppp | 234 | 45 | 262 | 51 | NA | NA | 40 | 36 | AB | Xaalppppp | 1.1134 | 44.5370 | 40.0833 | NA | 0.1962 | NA |
| ABalppaa | 9.9256 | 7862 | script | Packer | Xaalppaaap | 275 | 66 | 310 | 76 | NA | NA | 53 | 47 | AB | Xaalppaaa | 1.1418 | 60.5169 | 53.6659 | NA | 0.1962 | NA |
| ABalapaa | 9.4037 | 7827 | script | Packer | Xaalapaaaa | 323 | 118 | 353 | 125 | NA | NA | 52 | 46 | AB | Xaalapaaa | 1.1728 | 60.9851 | 53.9483 | NA | 0.1962 | NA |
| ABalapaa | 9.5992 | 7288 | script | Packer | Xaalapaaap | 266 | 61 | 300 | 71 | NA | NA | 52 | 46 | AB | Xaalapaaa | 1.1562 | 60.1203 | 53.1834 | NA | 0.1962 | NA |
| ABalapap | 9.8919 | 7407 | script | Packer | Xaalapappa | 247 | 51 | 277 | 59 | NA | NA | 45 | 41 | AB | Xaalapapp | 1.1400 | 51.2983 | 46.7385 | NA | 0.1962 | NA |
| ABalapap | 10.1720 | 7073 | script | Packer | Xaalapappa | 247 | 51 | 277 | 59 | NA | NA | 45 | 41 | AB | Xaalapapp | 1.1212 | 50.4560 | 45.9710 | NA | 0.1962 | NA |
| ABaraaaa | 9.8454 | 7011 | script | Packer | Xaaraaaaap | 278 | 70 | 313 | 80 | NA | NA | 52 | 46 | AB | Xaaraaaaa | 1.1391 | 59.2343 | 52.3996 | NA | 0.1962 | NA |
| ABalapap | 9.9903 | 7078 | script | Packer | Xaalapappp | 260 | 64 | 293 | 74 | NA | NA | 45 | 41 | AB | Xaalapapp | 1.1314 | 50.9146 | 46.3889 | NA | 0.1962 | NA |
| ABalppaa | 10.0097 | 6493 | script | Packer | Xaalppaapp | 273 | 66 | 308 | 76 | NA | NA | 51 | 45 | AB | Xaalppaap | 1.1248 | 57.3662 | 50.6173 | NA | 0.1962 | NA |
| ABalpaap | 10.5291 | 6946 | script | Packer | Xaalpaapaa | 251 | 61 | 283 | 71 | NA | NA | 40 | 36 | AB | Xaalpaapa | 1.1009 | 44.0376 | 39.6339 | NA | 0.1962 | NA |
| ABalppap | 9.8623 | 7619 | script | Packer | Xaalppapaa | 270 | 73 | 304 | 83 | NA | NA | 46 | 41 | AB | Xaalppapa | 1.1435 | 52.6001 | 46.8827 | NA | 0.1962 | NA |
| ABplppaa | 9.9722 | 7400 | script | Packer | Xaplppaapa | 280 | 73 | 316 | 85 | NA | NA | 50 | 45 | AB | Xaplppaap | 1.1353 | 56.7651 | 51.0886 | NA | 0.1962 | NA |
| ABplppaa | 10.2597 | 7738 | script | Packer | Xaplppaapp | 274 | 67 | 309 | 77 | NA | NA | 50 | 45 | AB | Xaplppaap | 1.1221 | 56.1042 | 50.4938 | NA | 0.1962 | NA |
| ABalaapa | 10.0097 | 6493 | script | Packer | Xaalaapapp | 274 | 64 | 309 | 74 | NA | NA | 52 | 46 | AB | Xaalaapap | 1.1248 | 58.4911 | 51.7421 | NA | 0.1962 | NA |
| ABplpapp | 10.7634 | 7320 | script | Packer | Xaplpappaa | 238 | 47 | 267 | 53 | NA | NA | 43 | 38 | AB | Xaplpappa | 1.0921 | 46.9610 | 41.5004 | NA | 0.1962 | NA |
| ABplpppp | 10.5519 | 8433 | script | Packer | Xaplppppaa | 259 | 63 | 292 | 72 | NA | NA | 46 | 40 | AB | Xaplppppa | 1.1117 | 51.1401 | 44.4697 | NA | 0.1962 | NA |
| ABplpppp | 10.5624 | 8080 | script | Packer | Xaplppppap | 259 | 63 | 292 | 72 | NA | NA | 46 | 40 | AB | Xaplppppa | 1.1086 | 50.9937 | 44.3424 | NA | 0.1962 | NA |
| ABplppap | 10.1155 | 7959 | script | Packer | Xaplppappp | 281 | 75 | 320 | 89 | NA | NA | 53 | 46 | AB | Xaplppapp | 1.1318 | 59.9869 | 52.0641 | NA | 0.1962 | NA |
| ABplppap | 9.8998 | 7504 | script | Packer | Xaplppappa | 274 | 68 | 309 | 77 | NA | NA | 53 | 46 | AB | Xaplppapp | 1.1403 | 60.4379 | 52.4555 | NA | 0.1962 | NA |
| ABpraapa | 10.2876 | 7327 | script | Packer | Xapraapapp | 254 | 56 | 285 | 63 | NA | NA | 48 | 42 | AB | Xapraapap | 1.1171 | 53.6228 | 46.9200 | NA | 0.1962 | NA |
| ABalpppp | 10.1227 | 7505 | script | Packer | Xaalppppap | 257 | 62 | 289 | 71 | NA | NA | 47 | 42 | AB | Xaalppppa | 1.1277 | 53.0029 | 47.3643 | NA | 0.1962 | NA |
| ABalpppp | 10.1498 | 7775 | script | Packer | Xaalppppaa | 263 | 68 | 296 | 78 | NA | NA | 47 | 42 | AB | Xaalppppa | 1.1284 | 53.0369 | 47.3947 | NA | 0.1962 | NA |
| ABprpppa | 10.3787 | 7877 | script | Packer | Xaprppppapa | 248 | 47 | 280 | 54 | NA | NA | 50 | 44 | AB | Xaprpppap | 1.1167 | 55.8373 | 49.1368 | NA | 0.1962 | NA |
| ABplapap | 10.2151 | 8925 | script | Packer | Xaplapappp | 270 | 77 | 304 | 88 | NA | NA | 46 | 40 | AB | Xaplapapp | 1.1335 | 52.1390 | 45.3383 | NA | 0.1962 | NA |
| ABalaapp | 9.8916 | 7606 | script | Packer | Xaalaapppa | 279 | 80 | 315 | 92 | NA | NA | 46 | 40 | AB | Xaalaappp | 1.1417 | 52.5169 | 45.6669 | NA | 0.1962 | NA |
| ABalaapp | 10.1274 | 7353 | script | Packer | Xaalaapppp | 256 | 57 | 288 | 65 | NA | NA | 46 | 40 | AB | Xaalaappp | 1.1262 | 51.8038 | 45.0468 | NA | 0.1962 | NA |
| ABalaapp | 10.1720 | 7073 | script | Packer | Xaalaapppp | 256 | 57 | 288 | 65 | NA | NA | 46 | 40 | AB | Xaalaappp | 1.1212 | 51.5772 | 44.8498 | NA | 0.1962 | NA |
| ABprapaa | 10.0829 | 7464 | script | Packer | Xaprapaaaa | 271 | 69 | 306 | 80 | NA | NA | 50 | 44 | AB | Xaprapaaa | 1.1296 | 56.4799 | 49.7023 | NA | 0.1962 | NA |
| ABprapaa | 10.2533 | 7523 | script | Packer | Xaprapaaap | 268 | 66 | 302 | 76 | NA | NA | 50 | 44 | AB | Xaprapaaa | 1.1207 | 56.0334 | 49.3094 | NA | 0.1962 | NA |
| ABalpapp | 10.4733 | 6429 | script | Packer | Xaalpappap | 242 | 51 | 272 | 58 | NA | NA | 44 | 38 | AB | Xaalpappa | 1.0990 | 48.3574 | 41.7632 | NA | 0.1962 | NA |
| ABalpapp | 10.1597 | 7976 | script | Packer | Xaalpappaa | 259 | 68 | 291 | 77 | NA | NA | 44 | 38 | AB | Xaalpappa | 1.1295 | 49.6980 | 42.9210 | NA | 0.1962 | NA |
| ABplpppa | 10.3787 | 7877 | script | Packer | Xaplpppapa | 247 | 47 | 278 | 53 | NA | NA | 49 | 43 | AB | Xaplpppap | 1.1167 | 54.7205 | 48.0200 | NA | 0.1962 | NA |
| ABprpapa | 10.2182 | 8062 | script | Packer | Xaprpapaaa | 270 | 68 | 304 | 78 | NA | NA | 52 | 46 | AB | Xaprpapaa | 1.1269 | 58.6005 | 51.8389 | NA | 0.1962 | NA |
| ABprpapa | 10.1385 | 7653 | script | Packer | Xaprpapaap | 270 | 68 | 304 | 78 | NA | NA | 52 | 46 | AB | Xaprpapaa | 1.1281 | 58.6599 | 51.8915 | NA | 0.1962 | NA |

| | | | | | | | | | | | | | | | | | | | | | |
|---|---|---|---|---|---|---|---|---|---|---|---|---|---|---|---|---|---|---|---|---|---|
| ABprppaa | 10.2790 | 7719 | script | Packer | Xaprppaaaa | 264 | 54 | 298 | 61 | NA | NA | 54 | 47 | AB | Xaprppaaa | 1.1209 | 60.5275 | 52.6814 | NA | 0.1962 | NA |
| ABaraapa | 10.0797 | 5783 | script | Packer | Xaaraapaaa | 268 | 69 | 302 | 79 | NA | NA | 47 | 42 | AB | Xaaraapaa | 1.1135 | 52.3345 | 46.7670 | NA | 0.1962 | NA |
| ABaraapa | 10.3066 | 7255 | script | Packer | Xaaraapaap | 256 | 57 | 288 | 65 | NA | NA | 47 | 42 | AB | Xaaraapaa | 1.1155 | 52.4283 | 46.8508 | NA | 0.1962 | NA |
| ABplpppp | 10.1514 | 6959 | script | Packer | Xaplpppppp | 260 | 61 | 293 | 70 | NA | NA | 49 | 43 | AB | Xaplppppp | 1.1214 | 54.9462 | 48.2181 | NA | 0.1962 | NA |
| ABplpppp | 10.4825 | 7404 | script | Packer | Xaplpppppa | 277 | 78 | 313 | 90 | NA | NA | 49 | 43 | AB | Xaplppppp | 1.1074 | 54.2606 | 47.6164 | NA | 0.1962 | NA |
| ABplppap | 10.2344 | 8384 | script | Packer | Xaplppapaa | 263 | 60 | 296 | 68 | NA | NA | 49 | 43 | AB | Xaplppapa | 1.1285 | 55.2960 | 48.5251 | NA | 0.1962 | NA |
| ABplppap | 10.1417 | 7431 | script | Packer | Xaplppapap | 278 | 75 | 315 | 88 | NA | NA | 49 | 43 | AB | Xaplppapa | 1.1260 | 55.1760 | 48.4197 | NA | 0.1962 | NA |
| ABarapap | 10.1597 | 7976 | script | Packer | Xaarapapaa | 261 | 67 | 294 | 76 | NA | NA | 46 | 40 | AB | Xaarapapa | 1.1295 | 51.9570 | 45.1800 | NA | 0.1962 | NA |
| ABarapap | 10.3947 | 6556 | script | Packer | Xaarapapap | 252 | 58 | 283 | 66 | NA | NA | 46 | 40 | AB | Xaarapapa | 1.1044 | 50.8029 | 44.1764 | NA | 0.1962 | NA |
| ABarapaa | 9.9706 | 5741 | script | Packer | Xaarapaapp | 247 | 50 | 277 | 57 | NA | NA | 44 | 39 | AB | Xaarapaap | 1.1191 | 49.2405 | 43.6450 | NA | 0.1962 | NA |
| ABarapaa | 10.0620 | 6464 | script | Packer | Xaarapaapa | 279 | 82 | 315 | 95 | NA | NA | 44 | 39 | AB | Xaarapaap | 1.1216 | 49.3510 | 43.7429 | NA | 0.1962 | NA |
| ABprpaap | 10.0276 | 7249 | script | Packer | Xaprpaappa | 259 | 62 | 292 | 72 | NA | NA | 46 | 41 | AB | Xaprpaapp | 1.1309 | 52.0192 | 46.3649 | NA | 0.1962 | NA |
| ABprpaap | 10.2240 | 7840 | script | Packer | Xaprpaappp | 257 | 60 | 290 | 70 | NA | NA | 46 | 41 | AB | Xaprpaapp | 1.1249 | 51.7438 | 46.1195 | NA | 0.1962 | NA |
| ABprpppa | 10.2193 | 7492 | script | Packer | Xaprpppaap | 269 | 68 | 304 | 79 | NA | NA | 49 | 44 | AB | Xaprpppaa | 1.1223 | 54.9914 | 49.3800 | NA | 0.1962 | NA |
| ABprpppa | 10.2821 | 7840 | script | Packer | Xaprpppaaa | 267 | 66 | 302 | 77 | NA | NA | 49 | 44 | AB | Xaprpppaa | 1.1217 | 54.9623 | 49.3539 | NA | 0.1962 | NA |
| ABarapap | 10.6826 | 7723 | script | Packer | Xaarapappp | 236 | 45 | 265 | 52 | NA | NA | 42 | 37 | AB | Xaarapapp | 1.0995 | 46.1804 | 40.6827 | NA | 0.1962 | NA |
| ABarapap | 10.0552 | 7039 | script | Packer | Xaarapappa | 288 | 97 | 326 | 113 | NA | NA | 42 | 37 | AB | Xaarapapp | 1.1274 | 47.3521 | 41.7149 | NA | 0.1962 | NA |
| ABarapaa | 10.1326 | 7216 | script | Packer | Xaarapaaaa | 271 | 67 | 306 | 77 | NA | NA | 53 | 46 | AB | Xaarapaaa | 1.1247 | 59.6084 | 51.7356 | NA | 0.1962 | NA |
| ABarapaa | 10.0594 | 6919 | script | Packer | Xaarapaaap | 279 | 75 | 316 | 87 | NA | NA | 53 | 46 | AB | Xaarapaaa | 1.1261 | 59.6833 | 51.8006 | NA | 0.1962 | NA |
| ABprpaap | 10.3265 | 7420 | script | Packer | Xaprpaapap | 266 | 66 | 300 | 76 | NA | NA | 50 | 44 | AB | Xaprpaapa | 1.1158 | 55.7914 | 49.0965 | NA | 0.1962 | NA |
| ABprpaap | 10.0720 | 7996 | script | Packer | Xaprpaapaa | 266 | 66 | 300 | 76 | NA | NA | 50 | 44 | AB | Xaprpaapa | 1.1346 | 56.7283 | 49.9209 | NA | 0.1962 | NA |
| ABplpaap | 10.0276 | 7249 | script | Packer | Xaplpaappa | 260 | 64 | 292 | 73 | NA | NA | 46 | 40 | AB | Xaplpaapp | 1.1309 | 52.0192 | 45.2341 | NA | 0.1962 | NA |
| ABalaapp | 9.9256 | 7862 | script | Packer | Xaalaappap | 273 | 67 | 308 | 77 | NA | NA | 53 | 47 | AB | Xaalaappa | 1.1418 | 60.5169 | 53.6659 | NA | 0.1962 | NA |
| ABalpapp | 10.0552 | 7039 | script | Packer | Xaalpapppa | 286 | 95 | 323 | 109 | NA | NA | 43 | 38 | AB | Xaalpappp | 1.1274 | 48.4795 | 42.8424 | NA | 0.1962 | NA |
| ABalpapp | 10.6826 | 7723 | script | Packer | Xaalpapppp | 235 | 44 | 264 | 50 | NA | NA | 43 | 38 | AB | Xaalpappp | 1.0995 | 47.2799 | 41.7823 | NA | 0.1962 | NA |
| ABpraaap | 10.2365 | 7691 | script | Packer | Xapraaappp | 312 | 74 | 347 | 79 | NA | NA | 52 | 46 | AB | Xapraaappp | 1.1230 | 58.3945 | 51.6567 | NA | 0.1962 | NA |
| ABpraaap | 10.2529 | 7947 | script | Packer | Xapraaappp | 323 | 85 | 352 | 85 | NA | NA | 52 | 46 | AB | Xapraaappp | 1.1241 | 58.4547 | 51.7100 | NA | 0.1962 | NA |
| ABalpppp | 10.2529 | 7947 | script | Packer | Xaalpppppa | 308 | 74 | 345 | 84 | NA | NA | 51 | 45 | AB | Xaalpppppp | 1.1241 | 57.3306 | 50.5858 | NA | 0.1962 | NA |
| ABalpppp | 10.2365 | 7691 | script | Packer | Xaalpppppp | 293 | 59 | 335 | 73 | NA | NA | 51 | 45 | AB | Xaalpppppp | 1.1230 | 57.2716 | 50.5337 | NA | 0.1962 | NA |
| C | 11.7241 | 7890 | manual | Tintori | Xppa | 67 | 17 | 66 | 18 | 18 | NA | 18 | NA | C | Xpp | 1.0508 | 18.9146 | NA | NA | 0.2264 | NA |
| Ca | 10.3854 | 7087 | script | Packer | Xppaa | 89 | 22 | 93 | 25 | 24 | 18 | 18 | 17 | C | Xppa | 1.1098 | 19.9761 | 18.8664 | 20.4201 | 0.2264 | 23.7854 |
| Cp | 10.3854 | 7087 | script | Packer | Xppap | 89 | 22 | 93 | 25 | 24 | 18 | 18 | 17 | C | Xppa | 1.1098 | 19.9761 | 18.8664 | 20.4201 | 0.2264 | 23.7854 |
| Ca | 10.8626 | 7020 | manual | Tintori | Xppaa | 89 | 22 | 93 | 25 | 24 | 18 | 18 | 17 | C | Xppa | 1.0846 | 19.5220 | 18.4374 | 19.9558 | 0.2264 | 23.7854 |
| Cp | 10.9511 | 7466 | manual | Tintori | Xppap | 89 | 22 | 93 | 25 | 24 | 18 | 18 | 17 | C | Xppa | 1.0839 | 19.5104 | 18.4265 | 19.9440 | 0.2264 | 23.7854 |
| Cpa | 10.1517 | 5863 | script | Packer | Xppapa | 118 | 29 | 126 | 32 | 29 | 24 | 25 | 22 | C | Xppap | 1.1104 | 27.7606 | 24.4293 | 26.6501 | 0.2264 | 31.0244 |
| Cpp | 10.1072 | 6819 | script | Packer | Xppapp | 121 | 32 | 130 | 36 | 32 | 24 | 25 | 22 | C | Xppap | 1.1225 | 28.0628 | 24.6952 | 26.9402 | 0.2264 | 31.0244 |
| Cap | 10.1072 | 6819 | script | Packer | Xppaap | 123 | 34 | 132 | 38 | 33 | 24 | 25 | 22 | C | Xppaa | 1.1225 | 28.0628 | 24.6952 | 27.0525 | 0.2264 | 31.1537 |
| Caa | 10.1517 | 5863 | script | Packer | Xppaaa | 119 | 30 | 127 | 33 | 29 | 24 | 25 | 22 | C | Xppaa | 1.1104 | 27.7606 | 24.4293 | 26.7612 | 0.2264 | 31.1537 |
| Capa | 10.2603 | 7262 | script | Packer | Xppaapa | 170 | 47 | 188 | 54 | 51 | 33 | 38 | 34 | C | Xppaap | 1.1181 | 42.4867 | 38.0144 | 37.3436 | 0.2264 | 43.1756 |
| Cpap | 10.3945 | 7084 | script | Packer | Xppapap | 153 | 35 | 167 | 39 | 35 | 29 | 32 | 29 | C | Xppapa | 1.1093 | 35.4969 | 32.1690 | 31.9472 | 0.2264 | 37.2293 |
| Cpaa | 10.0506 | 5875 | script | Packer | Xppapaa | 151 | 33 | 165 | 37 | 33 | 29 | 32 | 29 | C | Xppapa | 1.1161 | 35.7161 | 32.3677 | 32.1444 | 0.2264 | 37.2293 |
| Caap | 10.3945 | 7084 | script | Packer | Xppaaap | 160 | 41 | 175 | 46 | 42 | 29 | 33 | 30 | C | Xppaaa | 1.1093 | 36.6061 | 33.2783 | 32.3909 | 0.2264 | 37.7463 |
| Capp | 10.5051 | 7687 | script | Packer | Xppaapp | 164 | 41 | 180 | 47 | 43 | 33 | 38 | 34 | C | Xppaap | 1.1085 | 42.1228 | 37.6888 | 37.0237 | 0.2264 | 43.1756 |

| | | | | | | | | | | | | | | | | | | | | | |
|---|---|---|---|---|---|---|---|---|---|---|---|---|---|---|---|---|---|---|---|---|---|
| Caaa | 10.0506 | 5875 | script | Packer | Xppaaaa | 153 | 34 | 167 | 38 | 35 | 29 | 33 | 30 | C | Xppaaa | 1.1161 | 36.8322 | 33.4838 | 32.5909 | 0.2264 | 37.7463 |
| Cppp | 10.5051 | 7687 | script | Packer | Xppappp | 161 | 40 | 177 | 45 | 41 | 32 | 36 | 32 | C | Xppapp | 1.1085 | 39.9058 | 35.4718 | 35.9152 | 0.2264 | 41.8829 |
| Cppa | 10.2603 | 7262 | script | Packer | Xppappa | 168 | 47 | 185 | 53 | 49 | 32 | 36 | 32 | C | Xppapp | 1.1181 | 40.2505 | 35.7783 | 36.2255 | 0.2264 | 41.8829 |
| Caaaa | 10.0827 | 8055 | manual | Packer | Xppaaaaa | 197 | 44 | 218 | 49 | NA | NA | 38 | 34 | C | Xppaaaa | 1.1344 | 43.1082 | 38.5705 | NA | 0.2264 | NA |
| Caaap | 10.0827 | 8055 | manual | Packer | Xppaaaap | 194 | 41 | 215 | 46 | NA | NA | 38 | 34 | C | Xppaaaa | 1.1344 | 43.1082 | 38.5705 | NA | 0.2264 | NA |
| Cpaaa | 10.0827 | 8055 | manual | Packer | Xppapaaa | 197 | 46 | 219 | 52 | NA | NA | 37 | 33 | C | Xppapaa | 1.1344 | 41.9738 | 37.4361 | NA | 0.2264 | NA |
| Cpaap | 10.0827 | 8055 | manual | Packer | Xppapaap | 194 | 43 | 215 | 48 | NA | NA | 37 | 33 | C | Xppapaa | 1.1344 | 41.9738 | 37.4361 | NA | 0.2264 | NA |
| Cpapa | 10.2126 | 7323 | script | Packer | Xppapapa | 202 | 49 | 225 | 57 | NA | NA | 39 | 35 | C | Xppapap | 1.1212 | 43.7269 | 39.2421 | NA | 0.2264 | NA |
| Cpapp | 10.0396 | 8063 | script | Packer | Xppapapp | 203 | 50 | 226 | 58 | NA | NA | 39 | 35 | C | Xppapap | 1.1369 | 44.3401 | 39.7924 | NA | 0.2264 | NA |
| Caapa | 10.2227 | 8065 | script | Packer | Xppaaapa | 271 | 111 | 306 | 129 | NA | NA | 46 | 41 | C | Xppaaap | 1.1267 | 51.8287 | 46.1952 | NA | 0.2264 | NA |
| Cappp | 10.4320 | 7706 | script | Packer | Xppaappp | 215 | 51 | 240 | 59 | NA | NA | 47 | 41 | C | Xppaapp | 1.1125 | 52.2887 | 45.6135 | NA | 0.2264 | NA |
| Cappa | 10.3999 | 7778 | script | Packer | Xppaappa | 212 | 48 | 236 | 54 | NA | NA | 47 | 41 | C | Xppaapp | 1.1148 | 52.3964 | 45.7075 | NA | 0.2264 | NA |
| Capaa | 10.3391 | 7944 | script | Packer | Xppaapaa | 220 | 50 | 246 | 56 | NA | NA | 54 | 47 | C | Xppaapa | 1.1194 | 60.4479 | 52.6120 | NA | 0.2264 | NA |
| Capap | 10.3091 | 8248 | script | Packer | Xppaapap | 221 | 51 | 247 | 57 | NA | NA | 54 | 47 | C | Xppaapa | 1.1234 | 60.6623 | 52.7987 | NA | 0.2264 | NA |
| Cpppa | 10.3999 | 7778 | script | Packer | Xppapppa | 208 | 47 | 231 | 53 | NA | NA | 45 | 40 | C | Xppappp | 1.1148 | 50.1667 | 44.5926 | NA | 0.2264 | NA |
| Cpppp | 10.4320 | 7706 | script | Packer | Xppapppp | 212 | 51 | 236 | 58 | NA | NA | 45 | 40 | C | Xppappp | 1.1125 | 50.0636 | 44.5010 | NA | 0.2264 | NA |
| Cppap | 10.3091 | 8248 | script | Packer | Xppappap | 219 | 51 | 244 | 58 | NA | NA | 53 | 47 | C | Xppappa | 1.1234 | 59.5390 | 52.7987 | NA | 0.2264 | NA |
| Cppaa | 10.3391 | 7944 | script | Packer | Xppappaa | 218 | 50 | 243 | 57 | NA | NA | 53 | 47 | C | Xppappa | 1.1194 | 59.3285 | 52.6120 | NA | 0.2264 | NA |
| Caapp | 10.0396 | 8063 | script | Packer | Xppaaapp | 207 | 47 | 230 | 54 | NA | NA | 46 | 41 | C | Xppaaap | 1.1369 | 52.2985 | 46.6139 | NA | 0.2264 | NA |
| Cppppp | 10.2299 | 8025 | script | Packer | Xppappppp | 290 | 78 | 327 | 89 | NA | NA | 58 | 51 | C | Xppapppp | 1.1260 | 65.3082 | 57.4262 | NA | 0.2264 | NA |
| Cppppa | 10.0437 | 7882 | script | Packer | Xppappppa | 280 | 68 | 316 | 78 | NA | NA | 58 | 51 | C | Xppapppp | 1.1353 | 65.8446 | 57.8979 | NA | 0.2264 | NA |
| Cppapa | 9.9730 | 7516 | script | Packer | Xppappapa | 291 | 72 | 329 | 84 | NA | NA | 58 | 51 | C | Xppappap | 1.1362 | 65.9023 | 57.9485 | NA | 0.2264 | NA |
| Cppapp | 10.1106 | 7231 | script | Packer | Xppappapp | 301 | 82 | 341 | 96 | NA | NA | 58 | 51 | C | Xppappap | 1.1260 | 65.3106 | 57.4283 | NA | 0.2264 | NA |
| Cppaap | 10.0153 | 7316 | script | Packer | Xppappaap | 301 | 83 | 336 | 93 | NA | NA | 57 | 50 | C | Xppappaa | 1.1321 | 64.5315 | 56.6066 | NA | 0.2264 | NA |
| Cppaaa | 10.0141 | 7480 | script | Packer | Xppappaaa | 286 | 68 | 324 | 80 | NA | NA | 57 | 50 | C | Xppappaa | 1.1336 | 64.6156 | 56.6803 | NA | 0.2264 | NA |
| Capaap | 10.0153 | 7316 | script | Packer | Xppaapaap | 296 | 76 | 334 | 88 | NA | NA | 56 | 50 | C | Xppaapaa | 1.1321 | 63.3994 | 56.6066 | NA | 0.2264 | NA |
| Capaaa | 10.0141 | 7480 | script | Packer | Xppaapaaa | 287 | 67 | 326 | 80 | NA | NA | 56 | 50 | C | Xppaapaa | 1.1336 | 63.4820 | 56.6803 | NA | 0.2264 | NA |
| Cpppaa | 10.0440 | 7201 | script | Packer | Xppapppaa | 268 | 60 | 302 | 69 | NA | NA | 53 | 47 | C | Xppapppa | 1.1295 | 59.8637 | 53.0867 | NA | 0.2264 | NA |
| Capapp | 10.1106 | 7231 | script | Packer | Xppaapapp | 300 | 79 | 343 | 96 | NA | NA | 57 | 51 | C | Xppaapap | 1.1260 | 64.1845 | 57.4283 | NA | 0.2264 | NA |
| Capapa | 9.9730 | 7516 | script | Packer | Xppaapapa | 293 | 72 | 331 | 84 | NA | NA | 57 | 51 | C | Xppaapap | 1.1362 | 64.7660 | 57.9485 | NA | 0.2264 | NA |
| Capppp | 10.2299 | 8025 | script | Packer | Xppaapppp | 295 | 80 | 337 | 95 | NA | NA | 59 | 51 | C | Xppaappp | 1.1260 | 66.4342 | 57.4262 | NA | 0.2264 | NA |
| Capppa | 10.0437 | 7882 | script | Packer | Xppaapppa | 283 | 68 | 321 | 79 | NA | NA | 59 | 51 | C | Xppaappp | 1.1353 | 66.9799 | 57.8979 | NA | 0.2264 | NA |
| Cappaa | 10.0440 | 7201 | script | Packer | Xppaappaa | 275 | 63 | 310 | 72 | NA | NA | 54 | 48 | C | Xppaappa | 1.1295 | 60.9932 | 54.2162 | NA | 0.2264 | NA |
| Cappap | 10.2088 | 7810 | script | Packer | Xppaappap | 283 | 71 | 319 | 82 | NA | NA | 54 | 48 | C | Xppaappa | 1.1255 | 60.7750 | 54.0222 | NA | 0.2264 | NA |
| Cpppap | 10.2088 | 7810 | script | Packer | Xppapppap | 279 | 71 | 315 | 82 | NA | NA | 53 | 47 | C | Xppapppa | 1.1255 | 59.6495 | 52.8967 | NA | 0.2264 | NA |
| D | 10.8827 | 6802 | manual | Tintori | Xpppa | 107 | 34 | 113 | 38 | 33 | 26 | 25 | 23 | D | Xppp | 1.0816 | 27.0405 | 24.8773 | 27.7977 | 0.1190 | 29.1730 |
| Da | 10.1561 | 7712 | script | Packer | Xpppaa | 147 | 40 | 161 | 46 | 41 | 33 | 38 | 34 | D | Xpppa | 1.1276 | 42.8482 | 38.3379 | 37.2103 | 0.1190 | 37.4595 |
| Dp | 10.1561 | 7712 | script | Packer | Xpppap | 147 | 40 | 160 | 46 | 42 | 33 | 38 | 34 | D | Xpppa | 1.1276 | 42.8482 | 38.3379 | 37.2103 | 0.1190 | 37.4595 |
| Daa | 10.2084 | 7459 | script | Packer | Xpppaaa | 193 | 46 | 214 | 51 | 46 | 41 | 46 | 40 | D | Xpppaa | 1.1226 | 51.6392 | 44.9036 | 46.2507 | 0.1190 | 46.7676 |
| Dap | 10.2597 | 6476 | script | Packer | Xpppaap | 188 | 41 | 208 | 45 | 41 | 41 | 46 | 40 | D | Xpppaa | 1.1109 | 51.1003 | 44.4350 | 45.7681 | 0.1190 | 46.7676 |
| Dpp | 10.2597 | 6476 | script | Packer | Xpppapp | 187 | 40 | 207 | 45 | 40 | 42 | 46 | 40 | D | Xpppap | 1.1109 | 51.1003 | 44.4350 | 46.6568 | 0.1190 | 47.6757 |
| Dpa | 10.2084 | 7459 | script | Packer | Xpppapa | 192 | 45 | 212 | 50 | 45 | 42 | 46 | 40 | D | Xpppap | 1.1226 | 51.6392 | 44.9036 | 47.1488 | 0.1190 | 47.6757 |
| Daaa | 10.4729 | 9044 | manual | Packer | Xpppaaaa | 246 | 53 | 276 | 61 | NA | NA | 51 | 46 | D | Xpppaaaa | 1.1202 | 57.1319 | 51.5308 | NA | 0.1190 | NA |

| | | | | | | | | | | | | | | | | | | | | | |
|---|---|---|---|---|---|---|---|---|---|---|---|---|---|---|---|---|---|---|---|---|---|
| Dapa | 10.4729 | 9044 | manual | Packer | Xpppaapa | 242 | 54 | 271 | 62 | NA | NA | 45 | 41 | D | Xpppaap | 1.1202 | 50.4105 | 45.9296 | NA | 0.1190 | NA |
| Dpaa | 10.4729 | 9044 | manual | Packer | Xpppapaa | 248 | 56 | 279 | 65 | NA | NA | 50 | 45 | D | Xpppapa | 1.1202 | 56.0117 | 50.4105 | NA | 0.1190 | NA |
| Dppa | 10.4729 | 9044 | manual | Packer | Xpppappa | 244 | 57 | 274 | 65 | NA | NA | 45 | 40 | D | Xpppapp | 1.1202 | 50.4105 | 44.8094 | NA | 0.1190 | NA |
| Daap | 10.4865 | 7890 | script | Packer | Xpppaaap | 250 | 57 | 281 | 66 | NA | NA | 51 | 46 | D | Xpppaaa | 1.1111 | 56.6656 | 51.1101 | NA | 0.1190 | NA |
| Dppp | 10.6281 | 8599 | script | Packer | Xpppappp | 239 | 52 | 268 | 60 | NA | NA | 45 | 40 | D | Xpppapp | 1.1089 | 49.9024 | 44.3577 | NA | 0.1190 | NA |
| Dapp | 10.6281 | 8599 | script | Packer | Xpppaapp | 239 | 51 | 268 | 58 | NA | NA | 45 | 41 | D | Xpppaap | 1.1089 | 49.9024 | 45.4666 | NA | 0.1190 | NA |
| Dpap | 10.4865 | 7890 | script | Packer | Xpppapap | 250 | 58 | 281 | 67 | NA | NA | 50 | 45 | D | Xpppapa | 1.1111 | 55.5545 | 49.9990 | NA | 0.1190 | NA |
| Dappp | 10.2558 | 7555 | script | Packer | Xpppaappp | 330 | 91 | 362 | 92 | NA | NA | 58 | 51 | D | Xpppaapp | 1.1208 | 65.0064 | 57.1608 | NA | 0.1190 | NA |
| Dappa | 10.2800 | 7652 | script | Packer | Xpppaappa | 315 | 76 | 351 | 82 | NA | NA | 58 | 51 | D | Xpppaapp | 1.1203 | 64.9761 | 57.1341 | NA | 0.1190 | NA |
| Dpppa | 10.2800 | 7652 | script | Packer | Xpppapppa | 320 | 81 | 353 | 85 | NA | NA | 60 | 52 | D | Xpppappp | 1.1203 | 67.2166 | 58.2544 | NA | 0.1190 | NA |
| E | 11.5377 | 7996 | manual | Tintori | Xpap | 63 | 15 | 62 | 16 | 19 | NA | 14 | NA | E | Xpa | 1.0601 | 14.8407 | NA | NA | 0.2976 | NA |
| Ea | 10.2647 | 6578 | manual | Tintori | Xpapa | 101 | 38 | 106 | 43 | 38 | 19 | 16 | 15 | E | Xpap | 1.1116 | 17.7855 | 16.6739 | 21.0091 | 0.2976 | 26.9085 |
| Ep | 11.0381 | 7463 | manual | Tintori | Xpapp | 103 | 40 | 109 | 45 | 41 | 19 | 16 | 15 | E | Xpap | 1.0796 | 17.2738 | 16.1942 | 20.4047 | 0.2976 | 26.9085 |
| Eal | 10.2592 | 7534 | manual | Packer | Xpapal | 142 | 41 | 154 | 46 | 44 | 38 | 43 | 38 | E | Xpapa | 1.1204 | 48.1787 | 42.5765 | 43.0247 | 0.2976 | 54.6712 |
| Ear | 10.2592 | 7534 | manual | Packer | Xpapar | 143 | 42 | 155 | 47 | 44 | 38 | 43 | 38 | E | Xpapa | 1.1204 | 48.1787 | 42.5765 | 43.0247 | 0.2976 | 54.6712 |
| Epl | 10.2592 | 7534 | manual | Packer | Xpappl | 146 | 43 | 159 | 49 | 45 | 41 | 45 | 40 | E | Xpapp | 1.1204 | 50.4196 | 44.8174 | 45.4897 | 0.2976 | 57.8034 |
| Epr | 10.2592 | 7534 | manual | Packer | Xpappr | 147 | 44 | 160 | 50 | 45 | 41 | 45 | 40 | E | Xpapp | 1.1204 | 50.4196 | 44.8174 | 45.4897 | 0.2976 | 57.8034 |
| Eala | 10.2571 | 10428 | manual | Packer | Xpapala | 200 | 58 | 223 | 67 | NA | NA | 46 | 41 | E | Xpapal | 1.1408 | 52.4755 | 46.7716 | NA | 0.2976 | NA |
| Ealp | 10.2571 | 10428 | manual | Packer | Xpapalp | 202 | 60 | 225 | 69 | NA | NA | 46 | 41 | E | Xpapal | 1.1408 | 52.4755 | 46.7716 | NA | 0.2976 | NA |
| Eara | 10.2571 | 10428 | manual | Packer | Xpapara | 205 | 62 | 228 | 71 | NA | NA | 47 | 42 | E | Xpapar | 1.1408 | 53.6163 | 47.9124 | NA | 0.2976 | NA |
| Earp | 10.2571 | 10428 | manual | Packer | Xpaparp | 207 | 64 | 230 | 73 | NA | NA | 47 | 42 | E | Xpapar | 1.1408 | 53.6163 | 47.9124 | NA | 0.2976 | NA |
| Epla | 10.2571 | 10428 | manual | Packer | Xpappla | 208 | 62 | 231 | 71 | NA | NA | 49 | 43 | E | Xpappl | 1.1408 | 55.8978 | 49.0532 | NA | 0.2976 | NA |
| Eplp | 10.2571 | 10428 | manual | Packer | Xpapplp | 212 | 66 | 236 | 75 | NA | NA | 49 | 43 | E | Xpappl | 1.1408 | 55.8978 | 49.0532 | NA | 0.2976 | NA |
| Epra | 10.2571 | 10428 | manual | Packer | Xpappra | 210 | 63 | 234 | 72 | NA | NA | 50 | 44 | E | Xpappr | 1.1408 | 57.0386 | 50.1940 | NA | 0.2976 | NA |
| Eprp | 10.2571 | 10428 | manual | Packer | Xpapprp | 216 | 69 | 241 | 79 | NA | NA | 50 | 44 | E | Xpappr | 1.1408 | 57.0386 | 50.1940 | NA | 0.2976 | NA |
| EMS | 11.8898 | 8571 | manual | Tintori | Xpa | 48 | NA | 45 | 14 | NA | NA | NA | NA | MS | Xp | 1.0483 | NA | NA | NA | 0.1667 | NA |
| MS | 11.1746 | 7698 | manual | Tintori | Xpaa | 62 | 14 | 61 | 15 | 18 | NA | 14 | NA | MS | Xpa | 1.0749 | 15.0480 | NA | NA | 0.1667 | NA |
| MSp | 10.4459 | 5337 | script | Packer | Xpaap | 82 | 20 | 84 | 22 | 21 | 18 | 15 | 14 | MS | Xpaa | 1.0887 | 16.3309 | 15.2422 | 19.3794 | 0.1667 | 21.3600 |
| MSa | 10.4459 | 5337 | script | Packer | Xpaaa | 82 | 20 | 84 | 21 | 21 | 18 | 15 | 14 | MS | Xpaa | 1.0887 | 16.3309 | 15.2422 | 19.3794 | 0.1667 | 21.3600 |
| MSa | 11.1083 | 7323 | manual | Tintori | Xpaaa | 82 | 20 | 84 | 21 | 21 | 18 | 15 | 14 | MS | Xpaa | 1.0751 | 16.1258 | 15.0507 | 19.1359 | 0.1667 | 21.3600 |
| MSp | 11.0159 | 7987 | manual | Tintori | Xpaap | 82 | 20 | 84 | 22 | 21 | 18 | 15 | 14 | MS | Xpaa | 1.0848 | 16.2720 | 15.1872 | 19.3094 | 0.1667 | 21.3600 |
| MSpa | 10.3952 | 5058 | script | Packer | Xpaapa | 106 | 24 | 112 | 26 | 25 | 21 | 22 | 20 | MS | Xpaap | 1.0880 | 23.9351 | 21.7592 | 22.9559 | 0.1667 | 25.3200 |
| MSpp | 10.5352 | 6978 | script | Packer | Xpaapp | 108 | 26 | 115 | 29 | 27 | 21 | 22 | 20 | MS | Xpaap | 1.1009 | 24.2199 | 22.0181 | 23.2291 | 0.1667 | 25.3200 |
| MSap | 10.5352 | 6978 | script | Packer | Xpaaap | 108 | 26 | 114 | 29 | 27 | 21 | 21 | 20 | MS | Xpaaa | 1.1009 | 23.1190 | 22.0181 | 23.2291 | 0.1667 | 25.3200 |
| MSaa | 10.3952 | 5058 | script | Packer | Xpaaaa | 105 | 23 | 112 | 26 | 25 | 21 | 21 | 20 | MS | Xpaaa | 1.0880 | 22.8471 | 21.7592 | 22.9559 | 0.1667 | 25.3200 |
| MSapp | 10.5859 | 6733 | script | Packer | Xpaaapp | 137 | 29 | 149 | 33 | 31 | 27 | 29 | 26 | MS | Xpaaap | 1.0960 | 31.7853 | 28.4972 | 29.3740 | 0.1667 | 32.1600 |
| MSapa | 10.6107 | 7028 | script | Packer | Xpaaapa | 137 | 29 | 148 | 32 | 30 | 27 | 29 | 26 | MS | Xpaaap | 1.0974 | 31.8253 | 28.5330 | 29.4109 | 0.1667 | 32.1600 |
| MSaaa | 10.6775 | 6701 | script | Packer | Xpaaaaa | 133 | 28 | 144 | 30 | 29 | 25 | 26 | 23 | MS | Xpaaaa | 1.0910 | 28.3670 | 25.0939 | 27.0578 | 0.1667 | 29.7600 |
| MSaap | 10.7040 | 6999 | script | Packer | Xpaaaap | 133 | 28 | 144 | 31 | 29 | 25 | 26 | 23 | MS | Xpaaaa | 1.0924 | 28.4017 | 25.1246 | 27.0909 | 0.1667 | 29.7600 |
| MSpaa | 10.6775 | 6701 | script | Packer | Xpaapaa | 134 | 28 | 145 | 31 | 30 | 25 | 26 | 24 | MS | Xpaapa | 1.0910 | 28.3670 | 26.1850 | 26.8396 | 0.1667 | 29.5200 |
| MSpap | 10.7040 | 6999 | script | Packer | Xpaapap | 134 | 28 | 145 | 31 | 29 | 25 | 26 | 24 | MS | Xpaapa | 1.0924 | 28.4017 | 26.2170 | 26.8724 | 0.1667 | 29.5200 |
| MSppp | 10.5859 | 6733 | script | Packer | Xpaappp | 138 | 30 | 149 | 33 | 31 | 27 | 29 | 26 | MS | Xpaapp | 1.0960 | 31.7853 | 28.4972 | 29.3740 | 0.1667 | 32.1600 |
| MSppa | 10.6107 | 7028 | script | Packer | Xpaappa | 137 | 29 | 149 | 32 | 30 | 27 | 29 | 26 | MS | Xpaapp | 1.0974 | 31.8253 | 28.5330 | 29.4109 | 0.1667 | 32.1600 |

| | | | | | | | | | | | | | | | | | | | | | |
|---|---|---|---|---|---|---|---|---|---|---|---|---|---|---|---|---|---|---|---|---|---|
| MSpapp | 10.5710 | 7370 | script | Packer | Xpaapapp | 171 | 37 | 188 | 42 | 39 | 29 | 31 | 28 | MS | Xpaapap | 1.1024 | 34.1752 | 30.8680 | 31.8601 | 0.1667 | 34.6800 |
| MSpapa | 10.7442 | 7138 | script | Packer | Xpaapapa | 163 | 29 | 179 | 33 | 32 | 29 | 31 | 28 | MS | Xpaapap | 1.0915 | 33.8377 | 30.5631 | 31.5455 | 0.1667 | 34.6800 |
| MSpppa | 10.5172 | 7614 | script | Packer | Xpaapppa | 175 | 37 | 193 | 42 | 40 | 31 | 33 | 30 | MS | Xpaappp | 1.1073 | 36.5396 | 33.2179 | 33.8822 | 0.1667 | 36.7200 |
| MSpppp | 10.5837 | 6746 | script | Packer | Xpaapppp | 180 | 42 | 199 | 48 | 42 | 31 | 33 | 30 | MS | Xpaappp | 1.0963 | 36.1773 | 32.8885 | 33.5462 | 0.1667 | 36.7200 |
| MSappa | 10.5172 | 7614 | script | Packer | Xpaaappa | 177 | 40 | 195 | 44 | 42 | 31 | 33 | 29 | MS | Xpaaapp | 1.1073 | 36.5396 | 32.1106 | 34.2144 | 0.1667 | 37.0800 |
| MSappp | 10.5837 | 6746 | script | Packer | Xpaaappp | 180 | 43 | 199 | 49 | 44 | 31 | 33 | 29 | MS | Xpaaapp | 1.0963 | 36.1773 | 31.7922 | 33.8751 | 0.1667 | 37.0800 |
| MSaaaa | 10.6513 | 7322 | script | Packer | Xpaaaaaa | 162 | 29 | 178 | 33 | 32 | 29 | 30 | 28 | MS | Xpaaaaa | 1.0979 | 32.9359 | 30.7402 | 31.3989 | 0.1667 | 34.3200 |
| MSaaap | 10.4964 | 7510 | script | Packer | Xpaaaaap | 165 | 32 | 182 | 36 | 35 | 29 | 30 | 28 | MS | Xpaaaaa | 1.1075 | 33.2252 | 31.0101 | 31.6747 | 0.1667 | 34.3200 |
| MSapap | 10.4052 | 6929 | script | Packer | Xpaaapap | 170 | 33 | 188 | 38 | 36 | 30 | 32 | 29 | MS | Xpaaapa | 1.1073 | 35.4343 | 32.1123 | 33.4411 | 0.1667 | 36.2400 |
| MSapaa | 10.4868 | 7656 | script | Packer | Xpaaapaa | 171 | 34 | 188 | 38 | 37 | 30 | 32 | 29 | MS | Xpaaapa | 1.1092 | 35.4946 | 32.1670 | 33.4980 | 0.1667 | 36.2400 |
| MSppap | 10.4052 | 6929 | script | Packer | Xpaappap | 170 | 33 | 188 | 37 | 36 | 30 | 32 | 29 | MS | Xpaappa | 1.1073 | 35.4343 | 32.1123 | 33.4411 | 0.1667 | 36.2400 |
| MSppaa | 10.4868 | 7656 | script | Packer | Xpaappaa | 172 | 35 | 189 | 39 | 37 | 30 | 32 | 29 | MS | Xpaappa | 1.1092 | 35.4946 | 32.1670 | 33.4980 | 0.1667 | 36.2400 |
| MSaapa | 10.7442 | 7138 | script | Packer | Xpaaaapa | 162 | 29 | 178 | 33 | 32 | 29 | 31 | 28 | MS | Xpaaaap | 1.0915 | 33.8377 | 30.5631 | 31.7638 | 0.1667 | 34.9200 |
| MSaapp | 10.5710 | 7370 | script | Packer | Xpaaaapp | 170 | 37 | 187 | 42 | 39 | 29 | 31 | 28 | MS | Xpaaaap | 1.1024 | 34.1752 | 30.8680 | 32.0806 | 0.1667 | 34.9200 |
| MSpaaa | 10.6513 | 7322 | script | Packer | Xpaapaaa | 163 | 29 | 179 | 32 | 31 | 30 | 31 | 28 | MS | Xpaapaa | 1.0979 | 34.0337 | 30.7402 | 32.4967 | 0.1667 | 35.5200 |
| MSpaap | 10.4964 | 7510 | script | Packer | Xpaapaap | 169 | 35 | 186 | 39 | 38 | 30 | 31 | 28 | MS | Xpaapaa | 1.1075 | 34.3327 | 31.0101 | 32.7822 | 0.1667 | 35.5200 |
| MSpaaaa | 10.3919 | 6807 | script | Packer | Xpaapaaaa | 199 | 36 | 221 | 40 | NA | NA | 32 | 29 | MS | Xpaapaaa | 1.1069 | 35.4214 | 32.1006 | NA | 0.1667 | NA |
| MSpaaap | 10.3730 | 5895 | script | Packer | Xpaapaaap | 201 | 38 | 224 | 43 | NA | NA | 32 | 29 | MS | Xpaapaaa | 1.0989 | 35.1634 | 31.8668 | NA | 0.1667 | NA |
| MSpappa | 10.2472 | 7099 | script | Packer | Xpaapappa | 248 | 77 | 278 | 88 | NA | NA | 42 | 37 | MS | Xpaapapp | 1.1174 | 46.9288 | 41.3421 | NA | 0.1667 | NA |
| MSpappp | 10.6725 | 9005 | script | Packer | Xpaapappp | 234 | 63 | 262 | 72 | NA | NA | 42 | 37 | MS | Xpaapapp | 1.1094 | 46.5968 | 41.0496 | NA | 0.1667 | NA |
| MSpapap | 10.0168 | 7746 | script | Packer | Xpaapapap | 206 | 43 | 229 | 49 | NA | NA | 33 | 29 | MS | Xpaapapa | 1.1357 | 37.4773 | 32.9346 | NA | 0.1667 | NA |
| MSpapaa | 10.4785 | 8470 | script | Packer | Xpaapapaa | 199 | 36 | 221 | 40 | NA | NA | 33 | 29 | MS | Xpaapapa | 1.1159 | 36.8248 | 32.3611 | NA | 0.1667 | NA |
| MSpaapa | 10.4390 | 7803 | script | Packer | Xpaapaapa | 207 | 38 | 230 | 43 | NA | NA | 39 | 35 | MS | Xpaapaap | 1.1129 | 43.4042 | 38.9525 | NA | 0.1667 | NA |
| MSaaapp | 10.1559 | 6992 | script | Packer | Xpaaaaapp | 230 | 65 | 257 | 74 | NA | NA | 36 | 32 | MS | Xpaaaaap | 1.1214 | 40.3705 | 35.8849 | NA | 0.1667 | NA |
| MSaaapa | 10.4390 | 7803 | script | Packer | Xpaaaaapa | 199 | 34 | 222 | 38 | NA | NA | 36 | 32 | MS | Xpaaaaap | 1.1129 | 40.0654 | 35.6137 | NA | 0.1667 | NA |
| MSappaa | 10.6915 | 8401 | script | Packer | Xpaaappaa | 234 | 57 | 262 | 65 | NA | NA | 44 | 40 | MS | Xpaaappa | 1.1042 | 48.5860 | 44.1691 | NA | 0.1667 | NA |
| MSappap | 10.4963 | 8082 | script | Packer | Xpaaappap | 243 | 66 | 272 | 76 | NA | NA | 44 | 40 | MS | Xpaaappa | 1.1121 | 48.9306 | 44.4823 | NA | 0.1667 | NA |
| MSaappa | 10.3895 | 7277 | script | Packer | Xpaaaappa | 245 | 75 | 275 | 86 | NA | NA | 42 | 37 | MS | Xpaaaapp | 1.1112 | 46.6714 | 41.1153 | NA | 0.1667 | NA |
| MSapapp | 10.5820 | 8226 | script | Packer | Xpaaapapp | 222 | 52 | 248 | 58 | NA | NA | 38 | 33 | MS | Xpaaapap | 1.1086 | 42.1281 | 36.5850 | NA | 0.1667 | NA |
| MSppapp | 10.5820 | 8226 | script | Packer | Xpaappapp | 221 | 51 | 247 | 58 | NA | NA | 37 | 33 | MS | Xpaappap | 1.1086 | 41.0195 | 36.5850 | NA | 0.1667 | NA |
| MSaappp | 10.6725 | 9005 | script | Packer | Xpaaaappp | 236 | 66 | 265 | 76 | NA | NA | 42 | 37 | MS | Xpaaaapp | 1.1094 | 46.5968 | 41.0496 | NA | 0.1667 | NA |
| MSapppp | 10.5211 | 8774 | script | Packer | Xpaaapppp | 236 | 56 | 265 | 64 | NA | NA | 49 | 43 | MS | Xpaaappp | 1.1158 | 54.6746 | 47.9797 | NA | 0.1667 | NA |
| MSapppa | 10.4008 | 7752 | script | Packer | Xpaaapppa | 232 | 52 | 259 | 58 | NA | NA | 49 | 43 | MS | Xpaaappp | 1.1146 | 54.6135 | 47.9262 | NA | 0.1667 | NA |
| MSppaap | 10.2400 | 7032 | script | Packer | Xpaappaap | 233 | 61 | 261 | 70 | NA | NA | 39 | 35 | MS | Xpaappaa | 1.1171 | 43.5687 | 39.1002 | NA | 0.1667 | NA |
| MSaapaa | 10.4785 | 8470 | script | Packer | Xpaaaapaa | 198 | 36 | 220 | 40 | NA | NA | 33 | 29 | MS | Xpaaaapa | 1.1159 | 36.8248 | 32.3611 | NA | 0.1667 | NA |
| MSaapap | 10.0168 | 7746 | script | Packer | Xpaaaapap | 206 | 44 | 229 | 50 | NA | NA | 33 | 29 | MS | Xpaaaapa | 1.1357 | 37.4773 | 32.9346 | NA | 0.1667 | NA |
| MSapapa | 10.5459 | 8291 | script | Packer | Xpaaapapa | 213 | 43 | 238 | 49 | NA | NA | 38 | 33 | MS | Xpaaapap | 1.1110 | 42.2185 | 36.6635 | NA | 0.1667 | NA |
| MSppapa | 10.5459 | 8291 | script | Packer | Xpaappapa | 213 | 43 | 238 | 48 | NA | NA | 37 | 33 | MS | Xpaappap | 1.1110 | 41.1075 | 36.6635 | NA | 0.1667 | NA |
| MSppaaa | 10.5321 | 8103 | script | Packer | Xpaappaaa | 213 | 41 | 237 | 47 | NA | NA | 39 | 35 | MS | Xpaappaa | 1.1103 | 43.3028 | 38.8615 | NA | 0.1667 | NA |
| MSpppaa | 10.6915 | 8401 | script | Packer | Xpaapppaa | 232 | 57 | 260 | 65 | NA | NA | 42 | 37 | MS | Xpaapppa | 1.1042 | 46.3776 | 40.8564 | NA | 0.1667 | NA |
| MSpppap | 10.4963 | 8082 | script | Packer | Xpaapppap | 238 | 63 | 267 | 73 | NA | NA | 42 | 37 | MS | Xpaapppa | 1.1121 | 46.7065 | 41.1462 | NA | 0.1667 | NA |
| MSppppp | 10.5211 | 8774 | script | Packer | Xpaappppp | 236 | 56 | 265 | 65 | NA | NA | 48 | 42 | MS | Xpaapppp | 1.1158 | 53.5588 | 46.8639 | NA | 0.1667 | NA |
| MSppppa | 10.4008 | 7752 | script | Packer | Xpaappppa | 232 | 52 | 260 | 59 | NA | NA | 48 | 42 | MS | Xpaapppp | 1.1146 | 53.4990 | 46.8116 | NA | 0.1667 | NA |

| | | | | | | | | | | | | | | | | | | | | | |
|---|---|---|---|---|---|---|---|---|---|---|---|---|---|---|---|---|---|---|---|---|---|
| MSapaap | 10.5186 | 7344 | script | Packer | Xpaaapaap | 232 | 61 | 259 | 70 | NA | NA | 38 | 34 | MS | Xpaaapaa | 1.1050 | 41.9882 | 37.5684 | NA | 0.1667 | NA |
| MSapaaa | 10.5321 | 8103 | script | Packer | Xpaaapaaa | 213 | 42 | 238 | 48 | NA | NA | 38 | 34 | MS | Xpaaapaa | 1.1103 | 42.1925 | 37.7512 | NA | 0.1667 | NA |
| MSaaaap | 10.2774 | 7223 | script | Packer | Xpaaaaaap | 202 | 40 | 224 | 45 | NA | NA | 33 | 29 | MS | Xpaaaaaa | 1.1168 | 36.8543 | 32.3871 | NA | 0.1667 | NA |
| MSaaaaa | 10.3639 | 7244 | script | Packer | Xpaaaaaaa | 198 | 36 | 220 | 40 | NA | NA | 33 | 29 | MS | Xpaaaaaa | 1.1123 | 36.7062 | 32.2570 | NA | 0.1667 | NA |
| MSaaapa | 10.3519 | 6692 | script | Packer | Xpaaaaapap | 253 | 54 | 284 | 61 | NA | NA | 38 | 34 | MS | Xpaaaaapa | 1.1080 | 42.1033 | 37.6714 | NA | 0.1667 | NA |
| MSaaapa | 10.5981 | 7291 | script | Packer | Xpaaaaapaa | 247 | 48 | 277 | 54 | NA | NA | 38 | 34 | MS | Xpaaaaapa | 1.1004 | 41.8135 | 37.4120 | NA | 0.1667 | NA |
| MSaapap | 10.2596 | 7556 | script | Packer | Xpaaaapapp | 261 | 55 | 294 | 63 | NA | NA | 50 | 44 | MS | Xpaaaapap | 1.1206 | 56.0298 | 49.3062 | NA | 0.1667 | NA |
| MSaapap | 9.8467 | 7208 | script | Packer | Xpaaaapapa | 313 | 107 | 351 | 121 | NA | NA | 50 | 44 | MS | Xpaaaapap | 1.1408 | 57.0414 | 50.1964 | NA | 0.1667 | NA |
| MSpaapa | 10.1847 | 6808 | script | Packer | Xpaapaapaa | 282 | 75 | 318 | 86 | NA | NA | 43 | 38 | MS | Xpaapaapa | 1.1181 | 48.0794 | 42.4888 | NA | 0.1667 | NA |
| MSpaapa | 10.3519 | 6692 | script | Packer | Xpaapaapap | 258 | 51 | 290 | 58 | NA | NA | 43 | 38 | MS | Xpaapaapa | 1.1080 | 47.6432 | 42.1033 | NA | 0.1667 | NA |
| MSaaaap | 10.2231 | 6214 | script | Packer | Xpaaaaaapp | 266 | 64 | 300 | 74 | NA | NA | 45 | 40 | MS | Xpaaaaaap | 1.1102 | 49.9607 | 44.4095 | NA | 0.1667 | NA |
| MSaaaap | 10.3355 | 6715 | script | Packer | Xpaaaaaapa | 254 | 52 | 286 | 60 | NA | NA | 45 | 40 | MS | Xpaaaaaap | 1.1091 | 49.9085 | 44.3631 | NA | 0.1667 | NA |
| MSaaaaa | 10.4897 | 7470 | script | Packer | Xpaaaaaaap | 241 | 43 | 271 | 49 | NA | NA | 40 | 36 | MS | Xpaaaaaaa | 1.1075 | 44.3012 | 39.8711 | NA | 0.1667 | NA |
| MSpaaaa | 10.0122 | 6856 | script | Packer | Xpaapaaaaa | 282 | 83 | 320 | 97 | NA | NA | 40 | 36 | MS | Xpaapaaaa | 1.1282 | 45.1266 | 40.6140 | NA | 0.1667 | NA |
| MSpappp | 10.0074 | 7289 | script | Packer | Xpaapapppa | 301 | 67 | 340 | 79 | NA | NA | 72 | 63 | MS | Xpaapappp | 1.1323 | 81.5289 | 71.3378 | NA | 0.1667 | NA |
| MSpapap | 10.2596 | 7556 | script | Packer | Xpaapapapp | 261 | 55 | 293 | 63 | NA | NA | 49 | 43 | MS | Xpaapapap | 1.1206 | 54.9092 | 48.1856 | NA | 0.1667 | NA |
| MSpapap | 9.8467 | 7208 | script | Packer | Xpaapapapa | 300 | 94 | 340 | 110 | NA | NA | 49 | 43 | MS | Xpaapapap | 1.1408 | 55.9006 | 49.0556 | NA | 0.1667 | NA |
| MSpaaaa | 10.4627 | 7303 | script | Packer | Xpaapaaaap | 247 | 48 | 277 | 55 | NA | NA | 40 | 36 | MS | Xpaapaaaa | 1.1076 | 44.3020 | 39.8718 | NA | 0.1667 | NA |
| MSaapaa | 10.3985 | 8016 | script | Packer | Xpaaaapaaa | 250 | 52 | 281 | 60 | NA | NA | 40 | 36 | MS | Xpaaaapaa | 1.1168 | 44.6707 | 40.2036 | NA | 0.1667 | NA |
| MSaapaa | 10.4262 | 7855 | script | Packer | Xpaaaapaap | 245 | 47 | 275 | 54 | NA | NA | 40 | 36 | MS | Xpaaaapaa | 1.1140 | 44.5610 | 40.1049 | NA | 0.1667 | NA |
| MSpapaa | 10.4262 | 7855 | script | Packer | Xpaapapaap | 246 | 47 | 276 | 54 | NA | NA | 40 | 36 | MS | Xpaapapaa | 1.1140 | 44.5610 | 40.1049 | NA | 0.1667 | NA |
| MSpapaa | 10.3985 | 8016 | script | Packer | Xpaapapaaa | 251 | 52 | 282 | 60 | NA | NA | 40 | 36 | MS | Xpaapapaa | 1.1168 | 44.6707 | 40.2036 | NA | 0.1667 | NA |
| MSpaaap | 10.4022 | 7061 | script | Packer | Xpaapaaapp | 249 | 48 | 280 | 55 | NA | NA | 43 | 38 | MS | Xpaapaaap | 1.1087 | 47.6725 | 42.1292 | NA | 0.1667 | NA |
| MSpaaap | 9.9618 | 5931 | script | Packer | Xpaapaaapa | 275 | 74 | 310 | 84 | NA | NA | 43 | 38 | MS | Xpaapaaap | 1.1217 | 48.2331 | 42.6246 | NA | 0.1667 | NA |
| MSaappp | 10.0074 | 7289 | script | Packer | Xpaaaapppa | 309 | 73 | 348 | 85 | NA | NA | 76 | 66 | MS | Xpaaaappp | 1.1323 | 86.0583 | 74.7348 | NA | 0.1667 | NA |
| MSaaaaa | 10.3718 | 6806 | script | Packer | Xpaaaaaaap | 307 | 66 | 346 | 75 | NA | NA | 49 | 43 | MS | Xpaaaaaaap | 1.1080 | 54.2910 | 47.6431 | NA | 0.1667 | NA |
| P2 | 11.8403 | 7623 | manual | Tintori | Xpp | 50 | NA | 47 | 18 | NA | NA | NA | NA | P | Xp | 1.0436 | NA | NA | NA | NA | NA |
| P3 | 11.5398 | 7170 | manual | Tintori | Xppp | 73 | 23 | 73 | 25 | 26 | NA | 18 | NA | P | Xpp | 1.0535 | 18.9631 | NA | NA | NA | NA |
| P4 | 11.5361 | 7060 | manual | Tintori | Xpppp | 131 | 58 | 142 | 67 | 58 | 26 | 25 | 23 | P | Xppp | 1.0528 | 26.3189 | 24.2134 | 27.0559 | NA | NA |
